# Comparing methylation levels assayed in GC-rich regions with current and emerging methods

**DOI:** 10.1101/2023.09.06.556603

**Authors:** Dominic Guanzon, Jason P Ross, Chenkai Ma, Oliver Berry, Yi Jin Liew

## Abstract

DNA methylation is an epigenetic mechanism that regulates gene expression, and for mammals typically occurs on cytosines within CpG dinucleotides. A significant challenge for methylation detection methods is accurately measuring methylation levels within GC-rich regions such as gene promoters, as inaccuracies compromise downstream biological interpretation of the data. To address this challenge, we compared methylation levels assayed using four different methods: Enzymatic Methyl-seq (EM-seq), whole genome bisulphite sequencing (WGBS), Infinium arrays (Illumina MethylationEPIC, “EPIC”), and Oxford Nanopore Technologies nanopore sequencing (ONT) applied to human DNA. Overall, all methods produced comparable and consistent methylation readouts across the human genome. The flexibility offered by current gold standard WGBS in interrogating genome-wide cytosines is surpassed technically by both EM-seq and ONT, as their coverages and methylation readouts are less prone to GC bias. These advantages are tempered by increased laboratory time (EM-seq) and higher complexity (ONT). We further assess the strengths and weaknesses of each method, and provide recommendations in choosing the most appropriate methylation method for specific scientific questions or translational needs.

## Introduction

Attempts to characterise DNA methylation date back to the 1940s, where nucleic acids derived from calf thymus was separated using paper chromatography [1]. In that study, a small band “epi-cytosine” was identified as likely to be methylated cytosine, characterised by a methyl modification on carbon-5 of cytosine [2]. In the mammalian genome, this is typically followed by guanine and is referred to as a CpG site [3]. From 1970 onwards, DNA methylation research gained momentum through both advances in technical methods and demonstrations of the functional importance of methylation within biological systems. Specifically, DNA methylation can regulate gene expression [4], and that sodium bisulphite rapidly deaminates cytosine residues to uracil while methylated cytosine remains nonreactive [5].

The characterisation of methylation was further revolutionised by Frommer et al., who employed sodium bisulphite DNA conversion with strand-specific PCR amplification and subsequent DNA sequencing via the dideoxynucleotide chain-termination method [6]. This made available, for the first time, a reliable method to generate accurate methylation maps at the resolution of single DNA strands. The bisulphite-conversion method continued to underpin the later emergence of epigenomic technologies based on DNA microarray and next generation sequencing (NGS) technology [7]. Microarrays contain oligonucleotide probes on a solid support which produces a fluorescent signal upon hybridisation of suitably labelled DNA or RNA molecules [8]. Since 2008, Illumina have released the Infinium microarray design for profiling DNA methylation, with the latest of this technology being the Infinium MethylationEPIC (EPIC) BeadChip. The first version of EPIC assayed over 850,000 human CpG sites, while the improved second version covers over 935,000 sites. This array format has probes designed to hybridise with bisulphite-treated DNA derived from biological samples.

The next technological milestone for understanding genome-wide DNA methylation was the adaption of bisulphite-conversion to next-generation sequencing (NGS) workflows, allowing the sequencing of millions of DNA fragments in parallel [7]. In its early stages, NGS methylome studies commonly employed both complexity-reduction methods such as reduced representation bisulphite sequencing (RRBS) [9] and whole genome bisulphite sequencing (WGBS) methods [10, 11]. As the NGS technology matured and prices fell, WGBS became the predominant method adopted by methylome studies. RRBS covers approximately 6 million [12] while WGBS covers 28.2 million human CpG sites [13].

In general, there is good concordance between measurement of methylation levels derived from EPIC and WGBS data [14]. While WGBS offers more comprehensive methylome coverage, exceeding by an order of magnitude than EPIC microarrays [14], EPIC is popular for epigenome-wide association studies (EWAS). EWAS often considers phenotypes where the significant differences in DNA methylation between study groups may be subtle. In these instances, larger sample sizes, and hence library costs per sample, is paramount. To achieve roughly equivalent precision as EPIC measurements, very high coverages (mean > 30×) from WGBS is needed, adding to the cost pressures. Also, EPIC also requires less input DNA and offers easier bioinformatic analysis and interpretation [15].

While both WGBS and EPIC methods are highly effective, their reliance on bisulphite conversion introduces some limitations. Specifically, sodium bisulphite degrades DNA due to depyrimidination of unmethylated cytosines. These abasic sites are fragile and results in DNA strand breakage [16]. Consequently, WGBS typically yields low sequencing coverage across GC-rich regions. [17-19]. This is particularly problematic for characterising methylation state of key GC-rich functional regions of the genome, such as CpG islands. CpG islands are important for regulating gene expression and typically have abnormal methylation patterns in cancer [20-22]. Incorrectly interpreting the methylation state of GC-rich CpG islands has consequences for biological interpretation and identification of methylation biomarkers.

Two new laboratory techniques offer ways to characterise methylation levels without reliance on bisulphite conversion, and therefore without the bias it introduces: Enzymatic Methyl-seq (EM-seq) [18], and direct determination of methylation levels with third-generation DNA sequencing technologies [23, 24]. Instead of a chemical approach using sodium bisulphite, EM-seq uses TET2, an oxidation enhancer, and APOBEC for DNA conversion prior to sequencing with NGS. TET2 (with the oxidation enhancer) protects methylated cytosines through an oxidation cascade reaction; APOBEC converts unmethylated cytosines to uracil while protected methylated cytosines remain unchanged [18]. The less-destructive enzymatic conversion yields several advantages for EM-seq compared to WGBS, including more evenly spread dinucleotide distributions, higher coverage that is unbiased to GC content, and lower DNA input requirements [17, 18, 25].

The rapid development of third-generation sequencing techniques has opened new methods in detecting methylation status across longer loci. One example is the direct determination of cytosine methylation status sequenced using Oxford Nanopore Technologies (ONT) without prior conversion [23, 24]. The subtle difference in the electrical resistance generated when methylated and unmethylated cytosines passes through the pore is sufficient for machine learning techniques to produce a probabilistic score for the methylation status of each sequenced cytosine [26]. Like shorter reads, biases in methylation calls could result from sequence context, or due to imbalances in coverage. For the former, a systematic comparison of ONT methylation callers revealed that readout biases are largely attributable to the choice of methylation caller [27]. Furthermore, raw ONT base accuracy is slightly lower from GC-rich genomes [28] and from inverted duplicates [29], but this lower base quality has yet to be systematically linked to biases in methylation levels. For the latter, coverage from ONT reads are largely unaffected by local GC biases [30]; while methylation calls are mostly independent of coverage until it drops below 10× [27]. This motivated our efforts to investigate whether direct detection of methylated cytosines from ONT reads could offer a less-biased view of methylation, especially from GC-rich loci.

Although each of the technologies discussed above has its own strengths and weaknesses for detecting methylation, studies that conduct head-to-head comparisons on the same biological samples are lacking. Here, using matched human blood samples, we contrast genome-wide methylation readouts from EM-seq (enzymatically converted) against WGBS and EPIC microarrays (both bisulphite-converted). Given that the methylation states of CpG islands hold significant biological importance [20-22], our primary objective was to investigate whether enzymatic conversion or direct readouts yield more accurate methylation results in high GC-rich DNA—a known challenge for bisulphite conversion. To address this, we utilise a long loci (45S ribosomal DNA; ∼14 kb) which is extremely GC-rich, to compare the relative performances of EM-seq, WGBS and ONT. In addition to methodological advantages, we also considered practical factors such as cost and ease of use, ensuring that our findings provide contemporary and practical insights for researchers exploring these newer methods.

## Results

### Clinical specimens

Specimens were obtained from a previous diet study on the effect of fasting during a high protein, partial meal replacement program over a period of 16 weeks [31]. For this technical comparison work, DNA was extracted from whole bloods from two participants (WR025 and WR069) from two timepoints: visit 1 (*t* = 0, “V1”) and visit 9 (*t* = 16 weeks, “V9”) of the diet intervention, i.e., “WR025V1”, “WR025V9”, “WR069V1” and “WR069V9”. The same four samples were subsequently used in preparing libraries for EM-seq, WGBS, EPIC and ONT. Observed methylation patterns are likely similar to those from typical individuals in the population with no evidence of disease.

### Rarefaction of EM-seq and WGBS libraries to avoid analytical bias

Analysis of sequencing coverage revealed that EM-seq libraries had higher CpG coverage than sample-matched WGBS libraries. The modes of WGBS libraries ranged from 8–12×, while EM-seq had much higher modes of 10–40× (Fig. 1A). To reduce degree of coverage as a confounding factor for inter-library comparisons, raw reads—pre-trimmed, pre-mapped—were rarefied to match the coverage of the shallowest library (166 million reads; see Materials and Methods). This resulted in coverage patterns that were roughly equal among all libraries (8– 10×; Fig. 1A; Supplementary File S1). All downstream analyses were based on these rarefied libraries.

**Figure 1.**
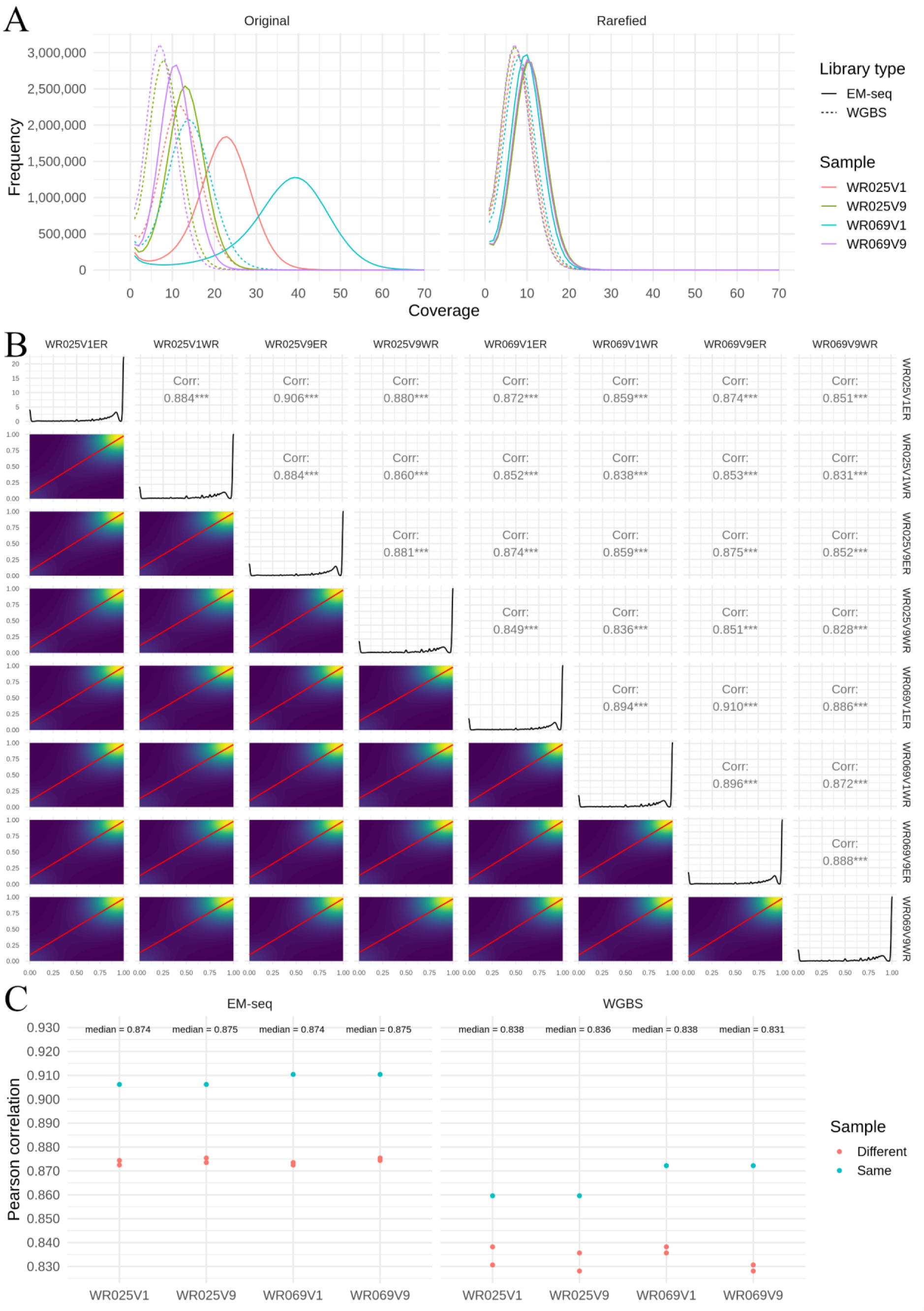
Methylation readouts from EM-seq libraries have better cross-sample correlations than WGBS libraries. (A) Original and rarefied coverages of EM-seq libraries (solid line) compared to WGBS libraries (dashed line), coloured by sample. (B) Pearson correlation was performed on rarefied data, comparing individual EM-seq (ending “-ER”) and WGBS (ending “-WR”) samples for 4 million randomly sampled CpG methylation beta values. The bottom left triangle contains pairwise sample comparisons represented as a 2D density plot. Red lines are lines of best fit, while the background shading indicates relative CpG density (blue: low; yellow: high). These sample comparisons are represented as Pearson correlations in the top right triangle, with *** indicating p-values < 0.001. The diagonal represents the distribution of CpG methylation beta values for individual samples. (C) Pearson correlation comparisons were separated by sample (x-axis) and grouped into EM-seq (left panel) and WGBS (right panel) library types. Comparison of samples from the same patient are coloured blue, while comparisons against different patients are coloured red.

### EM-seq libraries are more consistent and better covered in high GC regions than WGBS

All eight libraries had a similar bimodal distribution of methylation beta values, with heavy concentration of beta around 0 and 1 (0 indicates a fully unmethylated position, and 1 a fully methylated position; Fig. 1B), in line with previous observations [32]. To assess inter-library beta value correlations, we calculated Pearson correlation values from 4 million randomly sampled CpG sites. This analysis revealed significant positive correlations of methylation beta values between all samples, irrespective of the library preparation method (*r* = 0.826–0.906, all p values < 0.001; Fig. 1B). We further separated the Pearson correlations values by individuals and by library type. EM-seq libraries had significantly higher intra-method correlations than WGBS (mean *r* ± s.e.m., 0.885 ± 0.007 versus 0.844 ± 0.007, two-tailed *t*-test p < 0.01; Fig. 1C). Also, as expected, correlations of samples from the same patient (blue dots) were significantly higher than correlations with samples from different patients (red dots; 0.887 ± 0.011 versus 0.854 ± 0.007, two-tailed *t*-test *p* < 0.05; Fig. 1C).

Read GC content in EM-seq and WGBS libraries were analysed to identify whether libraries had differences in coverage in areas of high and low GC content. In line with previous studies [17, 18], EM-seq had elevated normalised coverage compared to WGBS in high GC regions (55–95 GC%) while WGBS had slightly higher normalised coverage in low GC regions (20– 35 GC%; Supplementary Fig. S1). In general, both libraries had low coverage in extreme low and high GC content regions (0 and 100 GC%; Supplementary Fig. S1).

### CpGs with significantly different betas have strand-specific and motif biases

We sought to identify CpG dinucleotides with significant beta value differences between EM-seq and WGBS libraries, and better understand factors associated with large discrepancies in estimated DNA methylation rates. From 28,704,358 CpGs analysed, only 124 CpGs (0.00043%) had significant differences in methylation betas across the library preparation methods (Supplementary File S2), indicating that both WGBS and EM-seq libraries overall have high CpG methylation beta concordance. We further analysed the discordance by clustering the beta values from the 124 CpGs in a heatmap with four other computed metrics representing strand-specific methylation, overall coverage, strand-specific coverage, and the immediate sequence context (3 bp either side of a CpG site). The clustering produced two distinct groups: one with significantly higher betas in WGBS than in EM-seq, and another vice versa (Fig. 2A).

**Figure 2.**
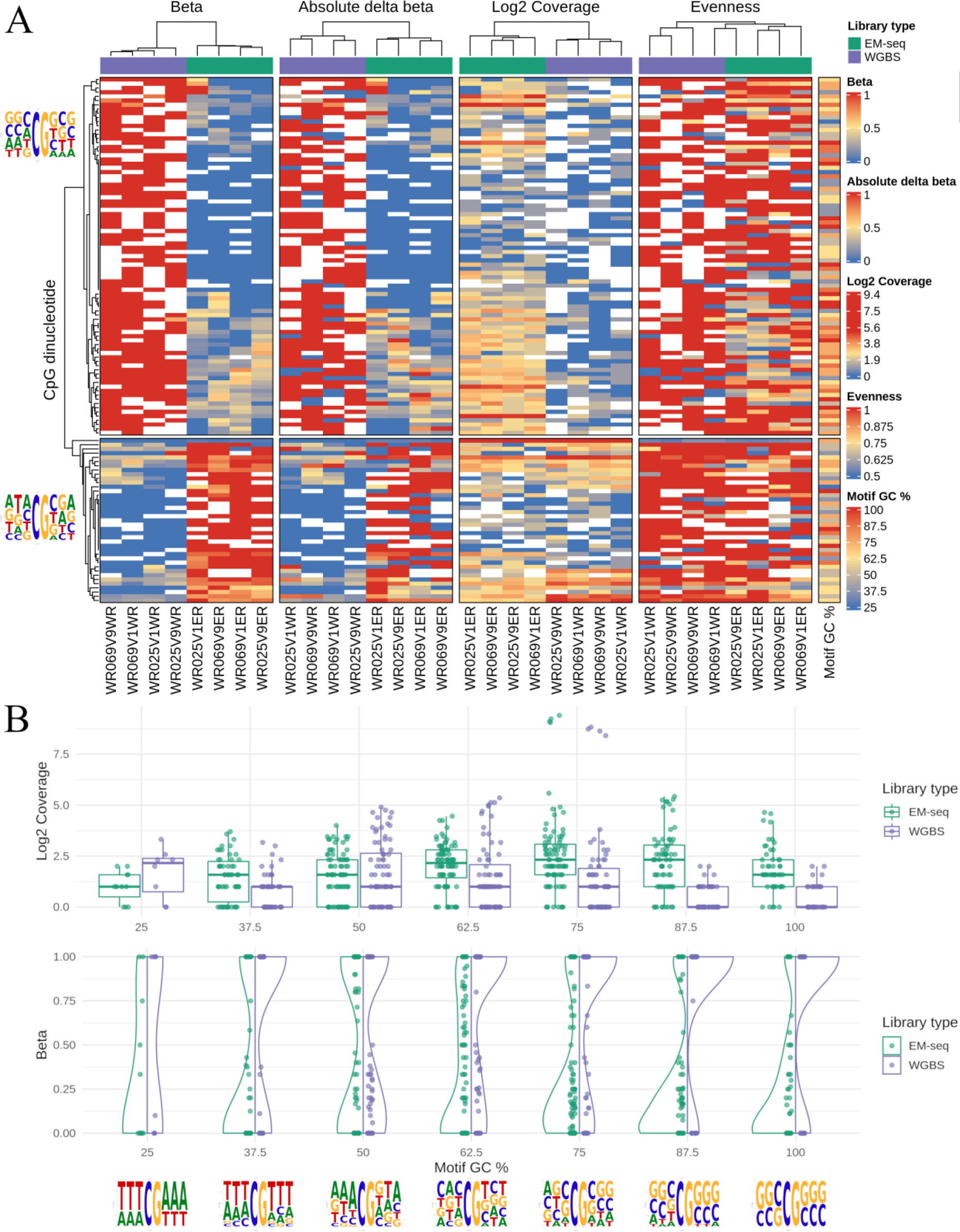
Strand-specific and motif biases associated with discordant methylation readouts. (A) Beta values from CpGs that were significantly different between EM-seq and WGBS libraries clustered into two groups in a heatmap. Top annotation bars represent library type, EM-seq (green) and WGBS (purple). Each column represents a different library; each row represents an individual CpG site. Dinucleotides with higher betas in WGBS are above, while those higher in EM-seq are below. Overall and strand-specific methylation levels are represented by “Beta” and “Absolute delta beta” respectively; overall and strand-specific coverages are represented by “Log2 coverage” and “Evenness” respectively. Sequence logos of 8 bp CpG motifs for both groups are placed beside the left dendrogram, while GC% of the motifs are on the rightmost column (“Motif GC%”). White boxes in the heatmap represent missing values. (B) Discordant CpGs were initially split by GC% of 8 bp motifs and subsequently by library type, EM-seq (green) and WGBS (purple). Individual bins were plotted against coverage (top panel) and beta (bottom panel). Sequence logos for each GC% bin is shown below the x-axis.

We sought to understand whether these discordant CpGs had strand-specific methylation readouts that departed from the assumed norm of being symmetrical (i.e., betas of Watson and Crick strands should be identical). We calculated two metrics for each CpG, one for beta and one for coverage, which we termed “absolute delta beta” and “evenness”, respectively. The former was calculated using the absolute differences in the strand-specific beta values from the Watson and Crick strands. We observed that absolute delta betas correlated with the methylation state at the CpG site, irrespective of library type (Fig. 2A). This previous finding is likely linked to the latter metric “evenness”, which was calculated using the ratio of the maximal single-strand coverage divided by combined coverage. If coverage was perfectly evenly split across Watson and Crick, then “evenness” is 0.5; if coverage was completely dominated by one strand (perfectly uneven), “evenness” is 1. The majority of differentially methylated CpG sites had coverage confined to a single strand, again irrespective of library type (Fig. 2A). Finally, coverages appeared to be lower for CpGs with significantly higher betas in WGBS (Fig. 2A). This suggests that the differential methylation readouts across both methods is an artefact resulting from coverage differences, where reads map asymmetrically to a single strand.

Next, we investigated whether the discordance in methylation readouts is linked to immediate sequence context. For each of the 124 CpGs, we derived an 8 bp motif comprising the CpG site with flanking 3 bp sequences (5’-NNN<u>CG</u>NNN-3’). Typically, CpGs with significantly higher betas in WGBS have motifs with high GC (Fig. 2A). In addition, these CpGs have a slight propensity to be flanked with “C” and “G” in the third and sixth positions of the motif. Conversely, CpGs with significantly higher betas in EM-seq have motifs with lower GC (Fig. 2A).

To further explore whether local GC content affects methylation betas and coverages, the 124 significant CpGs were split by the GC% value of their corresponding 8 bp CpG motifs (into seven bins) and by library preparation method (EM-seq or WGBS). EM-seq libraries had overall higher median coverage and lower methylation betas than WGBS, apart from coverage for motifs with GC% of 25% (Fig. 2B). Furthermore, these coverage differences were largest in motifs with high GC (62.5%–100%), and very low GC (25%). This relationship is reflected in the methylation betas, where lower coverage (for GC% of 62.5%–100%) resulted in more binary methylation betas of 0 and 1 for WGBS, compared to EM-seq which had a more even spread of methylation betas (Fig. 2B). If binary methylation betas were excluded, the remaining non-binary beta values were evenly distributed for EM-seq, compared to WGBS where values were mainly confined to betas of < 0.5. Interestingly, motifs with low GC (25%–50%) preferred to be flanked by either “A” or “T” 3 bp homopolymers. Finally, motifs with 50% GC percentage have the most even beta and coverage between library preparation methods, and strong preference for adenine in the -1 position (5’-NN**<u>A</u>**CGNNN-3’). Overall, there appears to be a relationship between coverage and motif GC% which contributes to differences in methylation readouts between EM-seq and WGBS.

### EM-seq readouts do not support previously reported TET2 biases

Echoing our motif-related observations, two recent studies have experimentally characterised the preference bias of TET enzymes for specific motifs surrounding methylated CpG dinucleotides [33, 34]. Specifically, TET2 (a component of EM-seq) has a binding preference for the 4-mer motif 5’-MCGW-3’, where M=A/C and W=A/T. Biases in TET2 activity are a concern, as its failure in catalysing 5-methylcytosine to 5-hydroxymethylcytosine will lead to APOBEC converting 5-methylcytosine to uracil (i.e., base will be misclassified as unmethylated).

We explored whether the set of 124 discordant CpGs had a significant enrichment/depletion of MCGW motifs, relative to the expected rates from the genome. For all biologically meaningful subdivisions of this dataset, the proportions of sequences with MCGW did not significantly depart from the norm (data not shown; available on GitHub). Subsequently, we extended the analysis to the full dataset, with the hopes of observing MCGW-driven differences in methylation properties that is exclusive to EM-seq and absent in WGBS. We were not able to find any meaningful differences in beta values, absolute delta beta, and evenness (Supplementary Fig. S2). Overall, our dataset indicates that the beta readouts from EM-seq do not support the previously reported TET2 preferences for MCGW.

### Methylation readouts from EPIC arrays equally comparable to EM-seq and WGBS

Using a principal components analysis, we observed that the variance in betas were predominantly due to library preparation method, not by sample ID (Supplementary Fig. S3). To simplify downstream analyses, we computed mean betas across all four samples from each method (EM-seq, WGBS and EPIC; circles in Supplementary Fig. S3). Readouts were largely similar: strong correlations were observed from all three between-method pairwise comparisons (*r* > 0.96, Figs. 3A, 3C and 3E). Whilst differences are slight, the short-read methods do agree with each other best (*r* = 0.97), followed by EPIC and EM-seq, and lastly EPIC and WGBS (both *r* = 0.96). The comparisons involving EPIC data had the greatest departures from the identity line (*y* = *x*), likely due to EPIC not being able to report perfectly unmethylated or methylated betas as 0 and 1 respectively due to autofluorescence during the scanning procedure (observed betas ranged from 0.0059–0.9933), and the addition of a default “stabilising constant” of 100 to the denominator in beta calculations (which prevents division by zero, but beta will always be < 1).

**Figure 3.**
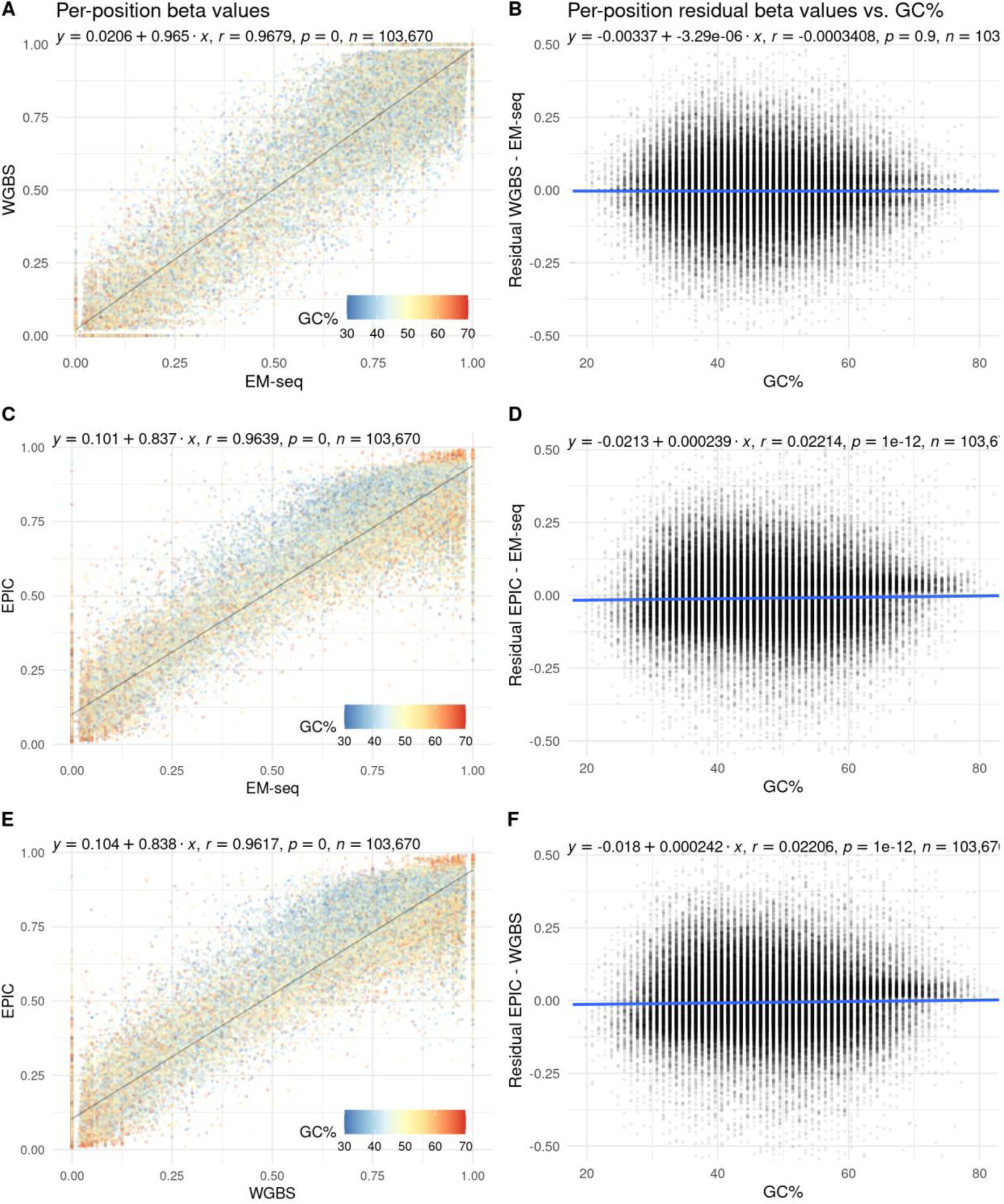
Per-position methylation levels assayed using EM-seq, WGBS and EPIC are well correlated and mostly independent of GC% context. (A) and (B) are for EM-seq vs. WGBS; (C) and (D) are for EPIC vs. EM-seq; while (E) and (F) are for EPIC vs. WGBS. Each point in the plots on the left (A, C and E) represent a single cytosine, and its position on the plot corresponds to the mean methylation level assayed using the method labelled on the axes. It is further coloured by the GC% of its local sequence context (±50 bp) to demonstrate context-dependent biases in methylation level measurements. For each point, the residual (differences in assayed methylation levels) was computed and plotted against the same GC% value on the right (B, D and F). All three trend lines are flat and close to origin. This implies that for well-covered cytosines, methylation readouts were almost independent of GC% context that are typical to these three methods (20–80%). At high GC values (> 75%), methylation levels appear to be higher in EPIC than either of EM-seq or WGBS (more points above the trendline in D and F; red points in the top right-hand corner of C and E).

Overall, GC% context had little-to-no influence on the eventual readouts of most positions common to these three methods. We defined local GC context as ±50 bp around the methylated cytosine. Between-method methylation readouts were mostly independent of typical GC contexts (20%–80%; Figs. 3B, 3D and 3F). At high GC (> 75%), EPIC overestimates methylation levels relative to both EM-seq and WGBS (Fig. 3C, 3D, 3E and 3F); or both short-read methods were underestimating actual methylation levels relative to EPIC. However, as there were few positions (*n* = 235) with GC% contexts of > 75%, broader conclusions could only be drawn from loci-specific experiments on a GC-rich loci.

EPIC probes were designed to target a subset of methylated CpGs in the human genome with cost-effectiveness in mind. As such, targeted positions are primarily located in biologically relevant and non-repetitive regions: mostly within gene bodies, or in promoter regions. Methylation readouts from all three methods were thus contrasted in a context-dependent manner (Fig. 4). Whilst correlation between all three methods were high in all six studied genomic contexts (*r* > 0.93), the short read methods (EM-seq and WGBS) were again in closer agreement than either was with EPIC (Fig. 4), like the earlier context-insensitive per-position analysis.

**Figure 4.**
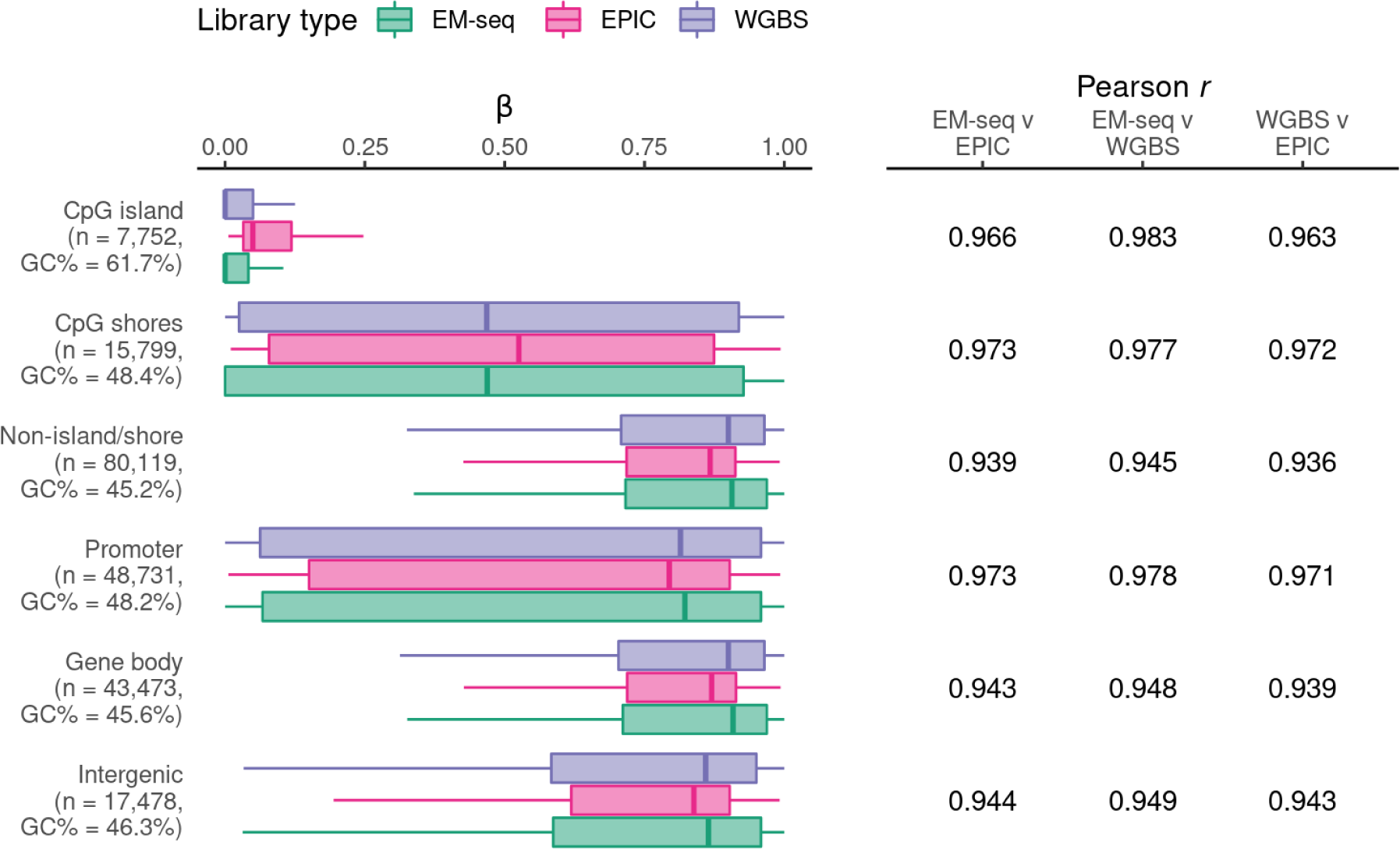
Methylation levels are well correlated in all biologically relevant genomic contexts across EM-seq, WGBS and EPIC. Positions were annotated and binned by their genomic contexts. The first three categories are exclusive—all positions are either in a CpG island, CpG shore, or neither. The next three categories are not exclusive as the human genome contains overlapping genes: a position could be in the gene body of the upstream gene, and in the promoter region of the downstream gene. Across all biologically relevant contexts with varying beta value and GC% ranges, the short read methods (EM-seq and WGBS) produced beta values that were in better agreement with each other than with EPIC.

### Direct methylation calls with ONT Cas9 more similar to EM-seq than WGBS in GC-rich loci

Few probes in the EPIC array are designed against highly GC-rich regions. To systematically benchmark the performance of EM-seq and WGBS in a longer GC-rich loci, the multicopy (∼300) human 45S rDNA gene was selected; this choice also allowed for ONT to be included into the technical comparison. The GC% content of this ∼14 kb locus is much higher than the genome mean (72% vs. 42%).

Our ONT Cas9 attempts achieved an estimated 350–480-fold increase in coverage along the 45S locus (Methods; Supplementary File S3). Each sample had approximately 1,500–2,500 reads mapping to the 45S locus (Supplementary Fig. S4), sufficient for downstream methylation level calling. A principal components analysis revealed that methylation levels showed more variance across methods than across biological replicates (Supplementary Fig. S5). The EM-seq replicates had the least inter-replicate variance; WGBS replicates had the highest. Means were separately calculated for the three methods and used in subsequent comparative purposes.

In terms of coverage, we observed more unique EM-seq reads mapping to the 45S locus than WGBS (Supplementary File S1). Initial sequencing depths were not a confounding factor, as we equalised the sequencing depths of all input files for this analysis. After conservatively removing cytosines with coverage values of < 50, the remaining 3,336 positions had comparable per-position coverages across all three methods (EM-seq mean: 947×; WGBS mean: 567×; ONT Cas9 mean: 608×). Coverage values from the short-read methods were inversely correlated with GC%, and EM-seq outperformed WGBS in terms of coverage at all measured GC% contexts (45%–95%, Fig. 5). ONT Cas9 coverages were mostly independent of GC% due to the length of the reads, and the DNA being read natively without conversion or PCR amplification (Fig. 5, Supplementary Fig. S6).

**Figure 5.**
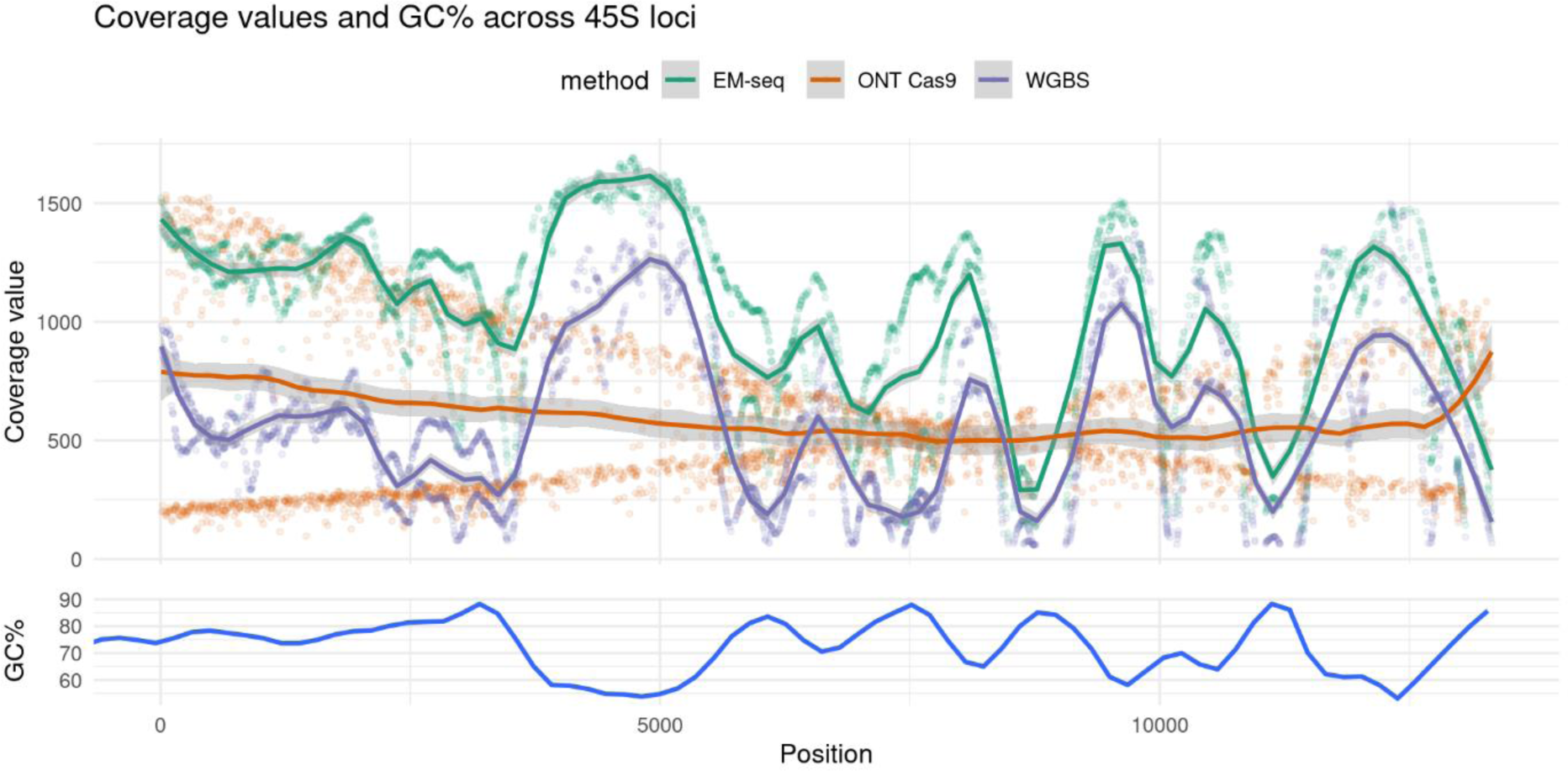
Per-position coverage values across the 45S locus. EM-seq and WGBS short reads have similar coverage profiles, with coverages inversely correlated with the GC% of the sequence context around the position (±50 bp). EM-seq however outperforms WGBS across the entire loci, with less stark dips in coverage at high GC% regions. ONT long reads have a “cross-shaped” coverage plot as the probe-guided Cas9 machinery makes blunt end cuts on one end (upstream/downstream of plotted region), while the other end is fragmented in a probabilistic manner.

In terms of beta values, EM-seq and WGBS reported similar betas when the local GC surrounding the measured cytosine was < 75% (Fig. 6B). Beyond that, WGBS progressively overestimated (or EM-seq underestimated) the observed betas, leading to the preponderance of red points above the line-of-best-fit in Fig. 6A, and the “hockey-stick” appearance of Fig. 6B. Despite this, these two Illumina sequencing-based methods still had a higher overall agreement with each other (*r* = 0.77; Fig. 6A), than either with ONT Cas9 (*r* = 0.54–0.58; Figs. 6C and 6E). The “hockey-stick” appearance in the residual plot of EM-seq vs. ONT Cas9 was still apparent, but less well-defined because of the poorer general agreement (Fig. 6D). At high GC (> 75%), ONT readouts were intermediate of the two short-read methods, but the intercepts and gradient values on the residual plots (flatter line closer to origin) indicate that ONT readouts were marginally closer to EM-seq than WGBS (Fig. 6D and 6F).

**Figure 6.**
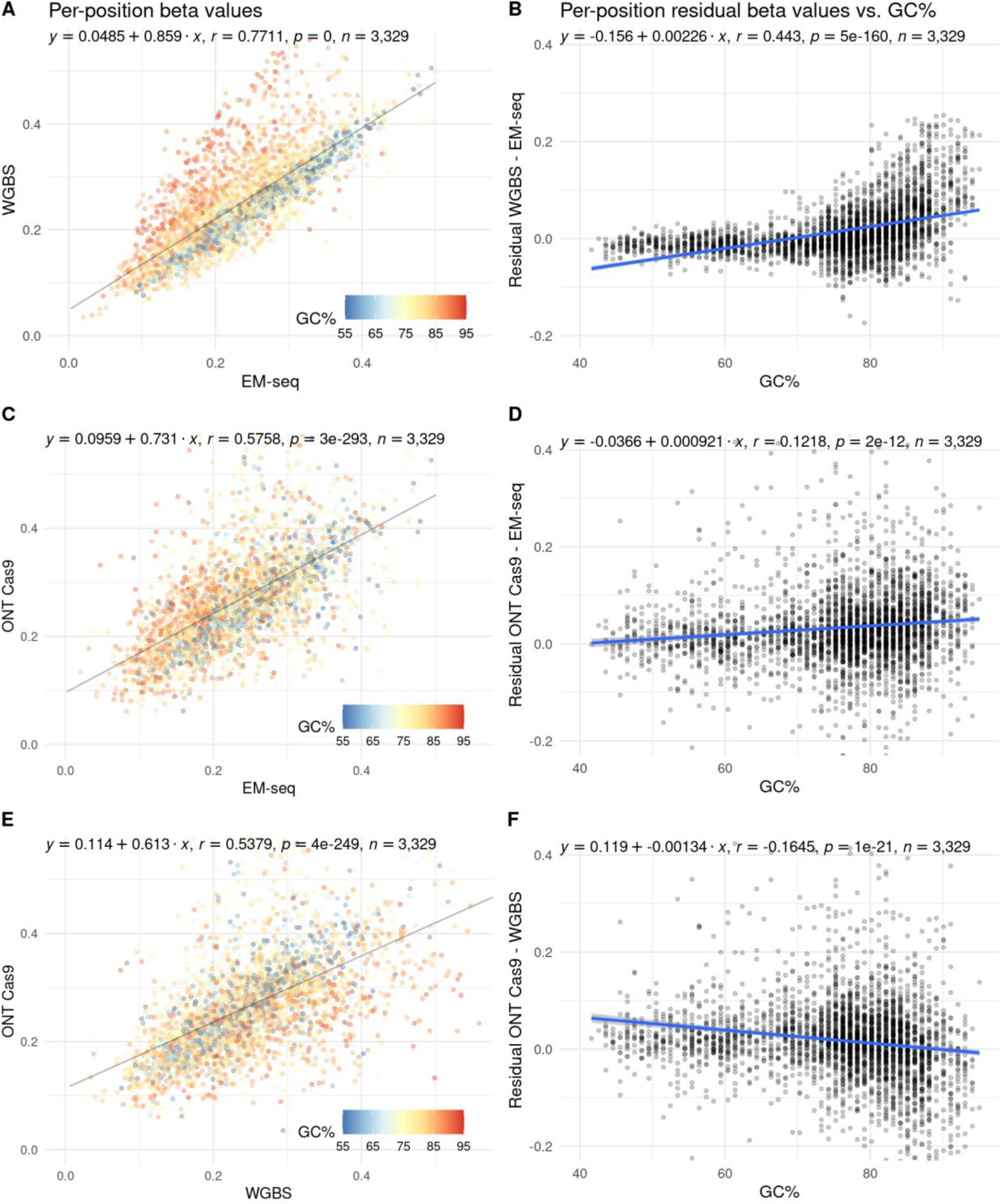
Pairwise comparisons of per-position methylation levels across the 45S locus. (A) and (B) are for EM-seq vs. WGBS; (C) and (D) are for ONT Cas9 vs. EM-seq; while (E) and (F) are for ONT Cas9 vs. WGBS. Each point in the plots on the left (A, C and E) represent a single cytosine, and its position on the plot corresponds to the mean betas assayed using the method labelled on the axes. It is further coloured by the GC% of its local sequence context (±50 bp) to demonstrate context-dependent biases in methylation level measurements. For each point, the residual (differences in assayed methylation levels) was computed and plotted against the same GC% value on the right (B, D and F). Overall trend lines indicate that at high GC levels (> 75%), WGBS overestimates methylation levels relative to EM-seq, while ONT readouts are intermediate of both EM-seq and WGBS. Axes for plots in the same column were equalised to facilitate visual comparisons.

## Discussion

### EM-seq vs. WGBS

As an alternative to bisulphite conversion of 5-methylcytosines, enzymatic conversion has several advantages: coverages that are less affected by GC context, lower DNA input requirements, and preservation of intact DNA strands [17, 18, 25]. In our hands, WGBS and EM-seq beta values were strongly correlated and highly concordant, which corroborates with previous reports [18, 35]. For GC-poor regions (< 40%), the coverage differences between EM-seq and WGBS were similar, as opposed to the reported 6-fold coverage difference for 10% GC regions [17, 18]. Conversely, we observed that EM-seq libraries have higher coverage in GC-rich regions compared to WGBS, supporting previous research [17, 18]. This finding holds biological significance, as the methylation of GC-rich regions, such as CpG islands and GC-rich repeat expansions, plays a pivotal role in cancer development [20-22] and monogenic disorders [36]. We recommend the use of EM-seq especially for studies focusing on GC-rich regions, as beta values estimated from higher coverages are more precise [37].

Whole genome analysis of 28,704,358 CpGs revealed that only 124 dinucleotides (0.00043%) had significantly different methylation betas comparing WGBS to EM-seq. We delved into the immediate sequence contexts surrounding these CpGs to elucidate possible explanations for the discordance. Strand bias has been observed in Illumina short read data, which is an important factor to consider for pathologies with strand-specific methylation patterns, such as the APC gene [38]. The majority of differentially methylated CpG sites had biased coverage to a specific strand, independent of library type. This bias leads to inter-strand methylation differences (absolute delta beta values) that closely approximate the overall beta value of the CpG dinucleotide. Overall, coverage differences are likely the cause of discrepancies in methylation calls between EM-seq and WGBS methods.

We hypothesised that discordant readouts could be the result of TET2, a key enzyme in EM-seq, which has reported preferences for converting specific motifs (5’-MCGW-3’) [33, 34]. From our data, TET2 did not seem to display the same motif preferences (Supplementary Fig. S2). New England Biolabs confirmed that use of the combination of EM-seq enzymes (TET2, Oxidation Enhancer and APOBEC) have been optimised to minimise sequencing bias (NEB, personal communication).

### EPIC vs. EM-seq vs. WGBS

Microarrays are a cost-effective tool for obtaining methylation information from larger sample sizes. Methylation readouts from EPIC have been shown to strongly correlate with both WGBS [14, 35] and EM-seq [35], with correlation coefficients all exceeding 0.85 from the cited studies. Our work builds upon these studies by confirming the strong correlations, and further framing methylation readouts across these three methods with local GC context.

As the EPIC protocol uses bisulphite-converted input DNA, like WGBS, we anticipated that beta values from EPIC would be more similar to WGBS instead of EM-seq. We observed that EPIC–EM-seq consistently had fractionally higher correlation values in all analysis than EPIC– WGBS. These observations were recapitulated in a similar analysis within the Supplementary section of Foox et al. [35], where EPIC–EM-seq had *r* = 0.853 and EPIC–WGBS had *r* = 0.852 (Swift WGBS was labelled as “MethylSeq”). While differences are minor, coverage is likely the reason driving this difference. As EM-seq has a more uniform coverage distribution across the genome than WGBS [17, 18, 35], beta values from the former are less “digital” (weaker clustering around common fractional values e.g., 0, 0.25, 0.33, 0.5, 1). This produces higher correlation values against “analogue” EPIC readouts (fluorescence readouts rarely cluster around common fractions).

For the subset of positions (*n* = 103,670) with sufficient data across all 12 datasets from three methods, betas were independent of window GC% (Fig. 3). However, for a small number of probes (*n* = 235) at high GC (> 75%), EPIC overestimates betas relative to both EM-seq and WGBS. This could be due to probes being cross-reactive, i.e., they hybridise to, and produce readouts from, off-targets elsewhere in the genome [14, 39]. Amongst the high GC probes, there were significantly more cross-reactive probes (*n* = 20 out of 235, 8.5%) than expected (*n* = 4,805 of 103,670, 4.6%; p < 0.01, Fisher’s exact test). To determine the broader biological implications for these observations, we extended the correlation analysis by grouping positions by their annotated genomic contexts. Due to the technical limitations leading to EPIC being less capable to report extreme beta values (of 0 and 1), the GC-rich, lowly methylated CpG islands had higher beta readouts in EPIC than both short-read methods. Overall, readouts remain strongly correlated across all genomic contexts in all three methods (*r* > 0.93), with EM-seq–WGBS having the strongest correlations.

### ONT vs. EM-seq vs. WGBS

The direct detection of methylated cytosines from ONT reads is still undergoing rapid improvements. This is not limited to only improvements in hardware including pore chemistry and library consumables, but also software such as modified base callers [26, 27]. Furthermore, recent variations in ONT sequencing e.g., Cas9-driven enrichment [40] and adaptive sequencing [41] provides alternative avenues to enrich for loci of interest, enabling higher coverages (and thus more accurate methylation readouts) for little-to-no additional effort.

By utilising an amplification-free, Cas9-driven enrichment of a GC-rich rDNA loci, we were able to compare methylation readouts from ONT direct detection against EM-seq and WGBS in this challenging context. In addition, the selection of a multicopy gene resulted in coverages that exceeded 500× for all three techniques, which increases accuracy of the methylation values. Our analysis revealed that EM-seq and WGBS had coverage values that were inversely correlated to local GC%, with WGBS having much stronger dips in coverage within extremely GC-rich regions (Fig. 5; Supplementary Figs. S6D, S6F). Of importance, we observed that ONT coverages remain unaffected by the same GC context bias, due to the long-read nature of this technology (Fig. 5). Previous literature also supports our findings, in the context of ONT sequencing of bacterial genomes with varying GC content [30]. Therefore, ONT is the optimal method for conducting studies involving genomes with large variations in GC contexts across the genome. In addition, shallower sequencing (< 30× coverage) due to cost considerations would still provide adequately informative methylation readouts in GC-rich contexts.

One challenge not commonly articulated about ONT modified base calling is that calls are based on confidence, rather than binary calls expected from sequencing converted DNA. For now, most methylation callers set confidence thresholds (e.g., 0-20% as unmethylated, 20-80% as undetermined, 80-100% as methylated) to generate beta values for comparative purposes. For this work, as the comparisons involved methods that generate binary calls, we used the default confidence thresholds of megalodon to generate ONT beta values. Correlations for ONT–EM-seq (*r* = 0.58) and ONT–WGBS (*r* = 0.54) were lower than EM-seq–WGBS (*r* = 0.77), most likely driven by greater technical commonalities between EM-seq and WGBS. Another reason would be that ONT readouts are the least biased by GC-context, followed by EM-seq and WGBS, with the latter having the most severe bias. This can be seen by uncertainties in measuring betas, quantified as the discrepancies in beta values (Figs. 6B, 6D, 6F), rising quickly in extremely GC-rich (> 75%) contexts. The overestimation of betas by WGBS in GC-rich contexts is in line with previous work that observed greater recovery of fully methylated fragments than fully unmethylated ones after bisulphite treatment [19].

It is important to note that the calling of modified bases from raw ONT signals is a rapidly developing field. Beta values differ based on methylation caller [27] and modified base models. Our current analysis used “remora”, a model trained purely on synthetic datasets (M.SssI-converted methylated DNA and PCR-amplified unmethylated DNA). Previously, when we used “rerio”, trained on a mix of WGBS data and synthetic datasets, ONT readouts behaved more like WGBS at extremely GC-rich contexts (ONT, personal communication; data available on GitHub). While future algorithmic refinement and improvement in ONT pore/chemistry could affect direct detection of modified bases, we remain confident that ONT methylation readouts would be less affected by GC-context biases than short-read methods.

### Practical considerations across all four methods

Ultimately, the choice of detection method is dependent on the biological question or translational needs. We focused on the relative performances of each method in GC-rich regions due to their relevance in cancer biology [20-22] and monogenic disorders [36]. We recommend selecting the method which is most cost-efficient and produces the highest quality data, especially when the question or need does not involve GC-rich regions. For brevity, this comparison is presented as a table (Table 1).

**Table 1.** Picking the right tool for the job. Practical considerations involved in all four methods, as well as their relative strengths and weaknesses.

| Criteria | EM-seq | WGBS | EPIC | ONT |
| --- | --- | --- | --- | --- |
| Flexibility in DNA conversion | NEB-only | Any bisulphite conversion kit |  | N/A |
| Flexibility in library construction | NEB-only | More options | Illumina-only | ONT-only |
| Flexibility in sequencing | Illumina-only | Depends on library type | Illumina-only | ONT-only |
| Experimental complexity | Well-established protocols which can be performed by trained scientists. |  |  | Protocols actively being developed and slightly more complex. |
| Data analysis complexity | Robust and mature packages/pipelines available |  |  | Pipelines are still in flux |
| Turnaround time (from DNA extracts) | 2-4 days |  | 2 days | 1-2 days (data is streamed) |
| Relative costs (per sample) | \$\$ | \$\$\$ | \$ | \$\$ (for ONT Cas9) |
| Strengths | Cheaper than WGBS. Coverage more evenly distributed across genome. Data quality better from | Easier to compare against publicly available data (most are WGBS/RRBS). Bisulphite conversion | Very cost effective for getting a subset of methylated and biologically relevant positions across more | Almost unbiased coverage regardless of context. Quickest turnaround time. Least affected by GC-context biases. |
|  | GC-rich loci than WGBS. | (without library building) cheaper than enzymatic conversion, better suited for translation into amplicon-based assays. | samples. Ideal for model organisms. |  |
| Weaknesses | Increased laboratory time than WGBS. Comparisons against existing data should consider readout divergences at GC-rich loci. | Coverage and methylation readouts biases very pronounced at GC-rich loci. | Not very practical for non-model organisms. Custom panels possible but less cost effective and less reliable. | Higher inputs required. Methylation data from whole genome possible, but more costly. Methylation calls are not binary, unlike bulk of existing data. Higher complexity in sequencing and in analysis. |

To conclude, both EM-seq and WGBS produces concordant methylation readouts, making either method a reliable choice for obtaining accurate data across the majority of cytosines in most genomes. While construction of EM-seq libraries is more time-consuming, they offer enhanced accuracies for readouts in GC-rich loci or genomes. On the other hand, EPIC presents a more cost-effective option, with the limitation of interrogating a smaller, predetermined set of positions in specific organisms. For researchers seeking unbiased coverage and rapid results, ONT emerges as a promising technology, particularly suitable for extremely GC-rich loci. However, it should be noted that ONT requires a slightly higher level of laboratory expertise and its analytical pipelines are still in the process of maturation.

## Methods

### Human ethics approval

Whole blood samples were collected from participants for a previous study that studied the dietary effects of fasting during a high protein, partial meal replacement program [31]. This trial was registered with the Australian New Zealand Clinical Trials Registry (http://www.anzctr.org.au; ACTRN12616000110482). Approval for this study is covered by CSIRO Health and Medical Human Research Ethics Committee (05/2015 and 2021_121_LR).

### Sample collection and DNA extraction

Four whole blood samples were obtained from two participants, WR025 and WR069 (both females, aged 32 and 28 respectively), at two collection timepoints: V1, at the start of the study and V9, 16 weeks into the study. Samples were labelled with participant ID and visit ID, i.e., “WR025V1”, “WR025V9”, “WR069V1” and “WR069V9”.

DNA was extracted from approximately 3 ml of whole blood using the Gentra Puregene Blood Kits (#158467; Qiagen, Hilden, Germany), following the “Whole Blood” subsection in the manufacturer-provided handbook. The optional RNase digestion step was carried out with the kit-provided RNase A solution. DNA quality and yields were assessed on a NanoDrop 1000 and Qubit 4 Fluorometer with Qubit dsDNA BR Assay kit (Thermo Fisher Scientific, Waltham, MA).

### Genome-wide/loci-specific methylation profiling with EM-seq and WGBS

For each sample, 1 µg of genomic DNA was pooled with 1:20 dilutions of unmethylated lambda (1 µl of 0.1 ng/µl) and methylated pUC19 control DNA (1 ul of 0.005 ng/µl) from the EM-seq kit (E7120L; NEB, Ipswich, MA). Volumes were made up to 50 ul with 0.1x TE buffer (Sigma-Aldrich, Burlington, MA). The pooled DNA samples were sheared to an average insert size of approximately 350 bp to 400 bp, using a Bioruptor (UCD400; Diagenode, Denville, NJ) on the high setting for 20-35 cycles (30 sec on, 30 sec off). Sheared DNA were run on a gel, and samples were subjected to additional shearing cycles if they were not sufficiently sheared. DNA concentrations were again quantified with a Qubit 4 Fluorometer (Thermo Fisher Scientific, Waltham, MA) to ensure DNA inputs for library creation were consistent across samples.

For EM-seq, 200 ng of sheared DNA was processed using the NEBNext Enzymatic Methyl-seq Kit (E7120; NEB, Ipswich, MA) following the manufacturer’s instructions for large insert libraries. For WGBS, 100 ng of sheared DNA was bisulphite-converted using the EZ DNA Methylation-Gold Kit (D5005; Zymo Research, Irvine, CA). Subsequently, the converted total DNA was processed using the Accel-NGS Methyl-Seq DNA Library Kit (#30024; Swift Biosciences, Ann Arbor, MI). EM-seq and WGBS libraries were quantified using the KAPA library quantification kit (KK4854; Roche Molecular Systems, Pleasanton, CA) and pooled in equimolar amounts. Pooled libraries were sequenced using the NovaSeq 6000 S4 2x150bp flowcell (Illumina, San Diego, CA) at the Ramaciotti Centre for Genomics (UNSW, Sydney, Australia), aiming for 30× coverage in perfectly pooled samples.

Analyses comparing WGBS against EM-seq in e.g., mapping rates, coverages, dinucleotide compositions, and per-position methylation levels are confounded by sequencing depth—for example, beta values are more accurate with greater sequencing depths. To remove this confounding effect, we equalised the sequencing depths of all short-read datasets used in this work to the shallowest one (166,282,895 reads). This rarefaction was carried out with a Python script (https://github.com/lyijin/common/blob/master/subsample_fastq.py), prior to read trimming.

The rarefied FASTQ files were processed using a self-written pipeline in Snakemake (https://github.com/lyijin/bismsmark) on the CSIRO High Performance Computing clusters. The pipeline depends on bismark v0.23.1 [42] and trim-galore v.0.6.7 (https://github.com/FelixKrueger/TrimGalore), while automatically applying tool-author-recommended command-line flags to deal with quirks associated with each method (https://github.com/FelixKrueger/Bismark/tree/master/Docs). For the WGBS datasets, the low-complexity bases added by the adaptase in the Accel-NGS kit necessitates the trimming of 10 bp from both ends of R1 and R2, and another 5 bp from off the 5’ end of R2 (i.e., 15 bp in total on the 5’ end of R2). For the EM-seq datasets, as the method-specific flags were added during manuscript preparation, they were treated as normal bisulphite-seq data: no special flags during trimming, but during extraction of methylation levels, methylation information in the first two bases on the 5’ end of R2 were discarded.

Genome-wide methylation levels were obtained by mapping the data (with default parameters) against the human GRCh38 patch 13 genome (https://www.ncbi.nlm.nih.gov/assembly/GCF_000001405.39/) with ALT and unplaced contigs removed. This was chosen as the updated Infinium MethylationEPIC array annotations (next section) is based on this version, allowing for comparisons that are free of annotation differences.

Loci-specific methylation levels were mapped against KY962518, a more modern 45S reference sequence produced with single-molecule sequencing [43], instead of U13369 that was pieced together from Sanger sequencing data from multiple labs [44]. The full KY962518 sequence contained a ∼13 kb transcribed region and a ∼32 kb intergenic spacer. As we were interested in methylation in the former region, we modified the sequence by placing the last 1 kb of the intergenic spacer (putative promoter region) in front of the ∼13 kb transcribed region, and discarded the remaining ∼31 kb of intergenic spacer. To deal with the slightly heterogenous rDNA reads arising from 45S genes that are not fully identical [45], we relaxed mapping parameters with --score-min L,0,-0.6 (bismark default is L,0,-0.2), which allowed reads with more mismatches to map to the 45S reference sequence. This resulted in higher coverages across the locus, and in most CpG dinucleotides, the methylation levels of cytosines on the Watson strand was closer to that on the Crick strand (more concordant methylation levels on both strands; data not shown).

Code written to parse bismark intermediate files into tabular form is at https://github.com/lyijin/cpgberus/tree/master/04_parse_bismark_covs, and code for EM-seq to WGBS comparisons analysed using bsseq (version 1.22.0) [46] and DSS R packages (version 2.34.0) [47] is at https://github.com/lyijin/cpgberus/tree/master/05_CpG_sequence_context with no smoothing or coverage cut-offs. Code that investigated potential MCGW-driven biases in EM-seq readouts is at https://github.com/lyijin/cpgberus/tree/master/13_check_mcgw_emseq_wgbs. Figures 1 and 2 were plotted using ggplot2 (version 3.3.3) [48], GGally (version 2.1.1) https://github.com/ggobi/ggally, ComplexHeatmap (version 2.2.0) [49] and motifStack (version 1.30.0) [50].

### Genome-wide methylation profiling with Illumina MethylationEPIC arrays

High molecular weight DNA (500–1,000 ng) was sent to the Australian Genome Research Facility (AGRF), Melbourne, Australia. DNA was subjected to bisulphite conversion, and methylation levels of over 850,000 sites were assayed with the Infinium MethylationEPIC BeadChip (Illumina, San Diego, CA).

Following receipt of data, the methylation levels associated with the four samples “WR025V1”, “WR025V9”, “WR069V1” and “WR069V9” were extracted using code documented in https://github.com/lyijin/cpgberus/tree/master/02_process_methepic_data. Raw data was subjected to noob correction [51] and then the beta values extracted and annotated with the Illumina manifest v1.0 B5 GRCh38 genome positions to maintain compatibility with WGBS and EM-seq read mapping. Some array features were discarded. These included 990 probes noted as high variability after a manufacturing change, 38 probes that did not have a GRCh38 genome location, 1,407 probes missing from the v1.0 B5 manifest and 36 pairs of probes mapping to the same GRCh38 coordinates. Illumina also supplied the genome coordinates as 0-based, so these were adjusted to 1-based coordinates.

### Three-way analysis of EPIC, EM-seq and WGBS data

The nature of the readouts was a key consideration in this three-way analysis: methylation readouts from EPIC were more “analogue”; while EM-seq and WGBS were more “digital” (e.g., methylation beta values of 0.50 are more common for the short-read methods than EPIC, as this results from having equal numbers of methylated and unmethylated reads). To reduce this “digital” effect, we picked positions that were covered ≥ 5 times in 3 of 4 EM-seq samples and similarly ≥ 5 times in 3 of 4 WGBS samples. This cut-off was stringent enough to minimise the “digital” effect, and fits with observations where gains in sensitivity is greatest going from 1× to 5× [52] yet lenient enough in not forcing all 4 samples to require the minimum 5× coverage that allowed for more positions for downstream analysis. Coverage values in WGBS samples were consistently lower than those for EM-seq: 27.6 million positions were covered ≥ 5 times in 3 of 4 EM-seq samples, while 12.4 million positions were covered ≥ 5 times in 3 of 4 WGBS samples. When further intersected with the > 850,000 positions from EPIC, we ended up with a common set of *n* = 103,670 positions with sufficient coverage for all downstream analysis. For these positions, the overall mean coverages from both short-read methods were comparable: 7.59 for EM-seq, 6.70 for WGBS. Our analysis proceeded with the assumption that this gap in coverage did not overly influence the per-position mean betas.

Code written to perform the three-way analysis of EPIC, EM-seq and WGBS data are available at https://github.com/lyijin/cpgberus/tree/master/14_methepic_vs_emseq_wgbs. Code used to annotate genomic context of probed positions in MethylationEPIC arrays utilised parsed databases generated with code at https://github.com/lyijin/cpgberus/tree/master/01_txdb.

### Loci-specific methylation profiling with ONT Cas9

CRISPR/Cas9 targeted sequencing was carried out using the Cas9 Sequencing Kit (SQK-CS9109; ONT, Oxford, UK) following the then-most recently available protocol (CAS_9106_v109_revC_16Sep2020) with modifications (Ramaciotti Centre for Genomics, UNSW Sydney, Australia). 1.25 µg of unsheared DNA from the same four samples, i.e., “WR025V1”, “WR025V9”, “WR069V1” and “WR069V9”, were used in this experiment. The dephosphorylating genomic DNA incubation time was increased to 20 minutes, and during the cleaving and dA-tailing DNA step, incubation was performed for 15 minutes.

To enable multiplexing of the four samples on one GridION flow cell, native barcoding was then performed (SQK-NBD114; ONT, Oxford, UK) as per the Cas9-targeted native barcoding protocol (Cas_native-v15). After barcoding and clean-up, all four samples were pooled in equal volume. Due to native barcoding been used, the adapter within the adapter ligation step was changed from the AMX adapter mix to AMII Adapter mix (supplied within SQK-NBD114).

Following manufacturer recommendations, probes were designed for this experiment using CHOPCHOP v3 [53] to target the conserved regions upstream and downstream of the 45S gene (Table 2). Notably, we disregarded ONT’s recommendation to pick probes with MM0 = 0, i.e., there should not be perfect matches to other parts of the genome. This is because the protocol assumes that experimenters are dealing with single-copy genes, with an MM0 of 0 implying no off-target events. In our case, the GRCh38 genome contained ∼10 copies of 45S of varying lengths across 5 chromosomes. The recommendation for MM1 and MM2 to both be 0 was still followed, to reduce likelihood of off-target (non-45S-targeting) events. Libraries constructed from the four separately barcoded DNA and common probes were then sequenced in a multiplexed manner on a single GridION flow cell (R9.4.1; ONT, Oxford, UK).

**Table 2.**
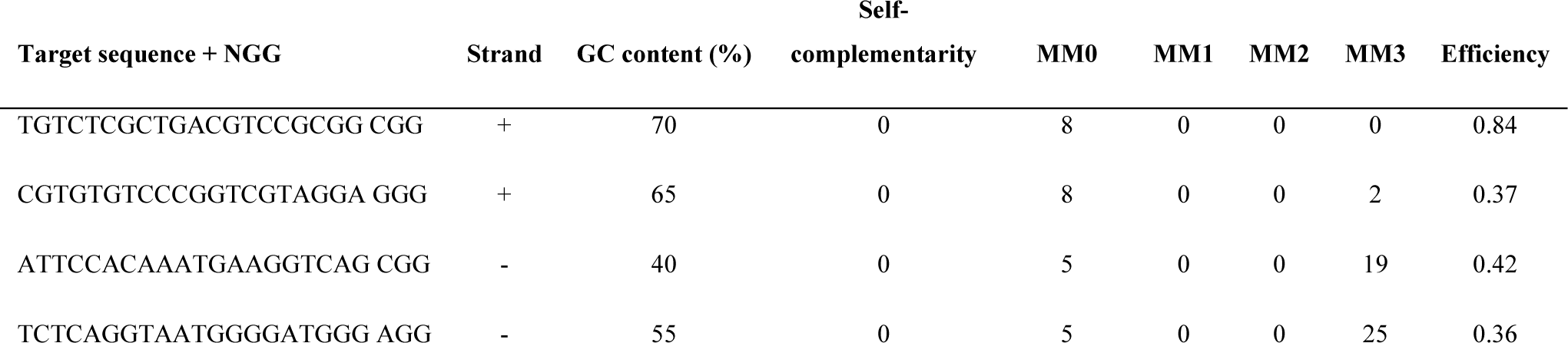
List of forward and reverse probes used in the ONT Cas9 experiment. The columns “strand” to “efficiency” are outputs from CHOPCHOP. “MM*n*” is the number of matches to other parts of the genome with *n* mismatches (i.e., MM0 implies perfect matches). ONT recommends for probes to have a GC% of 40–80%, self-complementarity score of 0, MM0 = 0, MM1 = 0, MM2 = 0 and MM3 ≤ 5. Efficiency scores are 0–1, higher values indicate higher likelihood of success.

Per-base methylation levels were directly called from PASS-quality ONT reads in the respective FAST5 files using an ONT-authored tool, megalodon v2.4.2 (https://github.com/nanoporetech/megalodon). This tool had two dependencies, guppy v5.0.14 (from https://community.nanoporetech.com/downloads, requires login) and remora v0.1.2 (https://github.com/nanoporetech/remora). The setting up of megalodon to utilise GPUs on the CSIRO cluster for faster calls, and the hacks needed to perform calls on a per-barcode basis (not supported by default), is documented at https://github.com/lyijin/cpgberus/tree/master/06_process_ont_data.

Due to the multi-kb nature of the reads, the 45S reference sequence (modified KY962518) used for the short WGBS and EM-seq was not appropriate in the mapping of these long reads. We constructed another modified KY962518 sequence where we transposed the last 11 kb of the sequence to the start of the sequence (i.e., the 12,345^th^ base from the 5’ end of this modified reference would be the 2,345^th^ bp in the one used for short reads, and 1,345^th^ in the original KY962518 sequence).

To avoid confusing readers, the choice of reference sequences has been masked in analyses and plots by making sure the transcribed regions of all reference sequences start at +1, i.e., treating the first 11 kb of the long-read reference as -11,000 to -1, and treating the first 1 kb of the short-read reference as -1,000 to -1.

Code written for the loci-specific analysis is available at https://github.com/lyijin/cpgberus/tree/master/15_ont_minimap2_coverage and https://github.com/lyijin/cpgberus/tree/master/16_loci_specific_three_way.

## Data access

Due to privacy concerns, sequence data in FASTQ/FAST5 format is not available. Intermediate tab-delimited files with per-position beta values and genomes used for mapping are available at https://data.csiro.au/collection/csiro:58492.

Code for this project is available at https://github.com/lyijin/cpgberus.

## Acknowledgements

We thank Jane Bowen and colleagues for sample collection; Thu Ho for assistance in DNA extraction; the Australian Genome Research Facility (Melbourne, Australia) for conducting the microarray experiments; Ramaciotti Centre for Genomics (UNSW Sydney, Australia) for short-read library sequencing; Tonia Russell for carrying out the ONT Cas9 work; and Bioplatforms Australia and Environomics FSP (CSIRO) for funding.

## Author contributions

Y.J.L designed the study, with mentorship and advice from J.R. and O.B.; D.G., C.M. and Y.J.L. performed experiments; D.G., J.R., Y.J.L. analysed data and generated plots; D.G. and Y.J.L. wrote the initial manuscript; all authors edited and commented on the final manuscript.

## Supplementary Figures

**Supplementary Figure S1.**
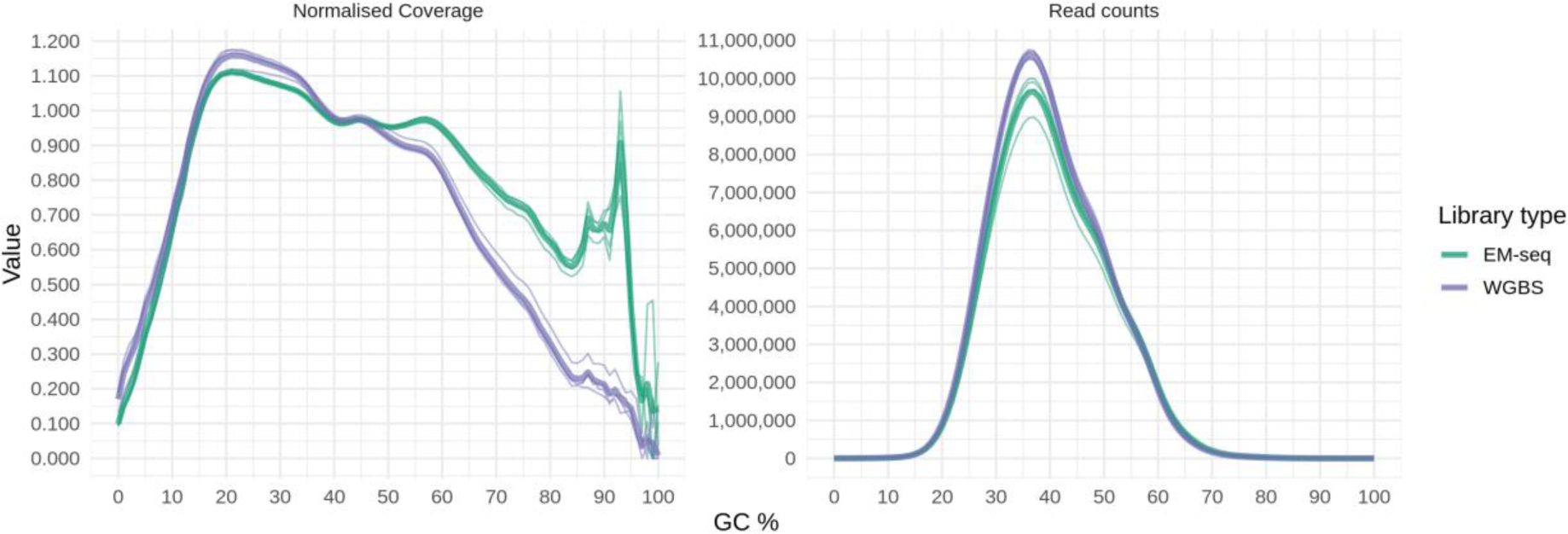
EM-seq libraries have higher coverage in GC-rich regions. Normalised coverage (left panel) and read counts (right panel), in EM-seq libraries (green) and WGBS libraries (blue) compared to GC% in 100 base pair bins. Thick lines and thin lines represent the average and individual samples, respectively.

**Supplementary Figure S2.**
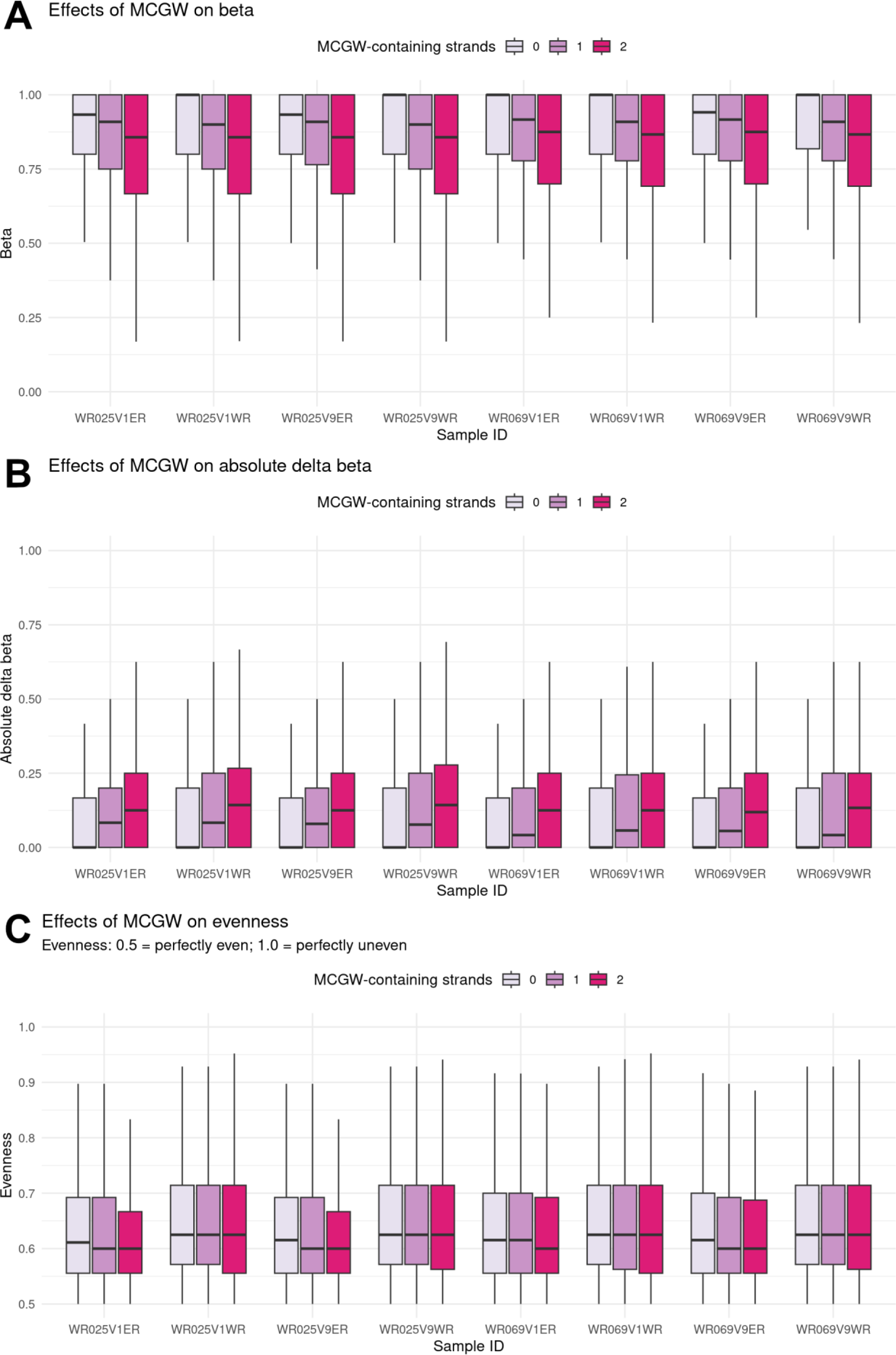
Methylation readouts binned by MCGW status for all eight samples. All CpGs with sufficient coverage (≥ 5; > 22 million dinucleotides for all samples) were categorised by MCGW-containing strands. As MCGW is not a palindromic motif, out of the overall 16 NCGN possibilities, 9 contexts have neither strand with MCGW, 6 have a single strand with MCGW, and 1 where both strands are MCGW (ACGT/ACGT). If EM-seq libraries had biased beta readouts linked with MCGW status, the distribution of (A) beta values, (B) absolute delta beta values and (C) evenness would be different in samples with “-ER” (EM-seq libraries) than those with “-WR” (WGBS). In all three cases, MCGW status has little-to-no effect on those values.

**Supplementary Figure S3.**
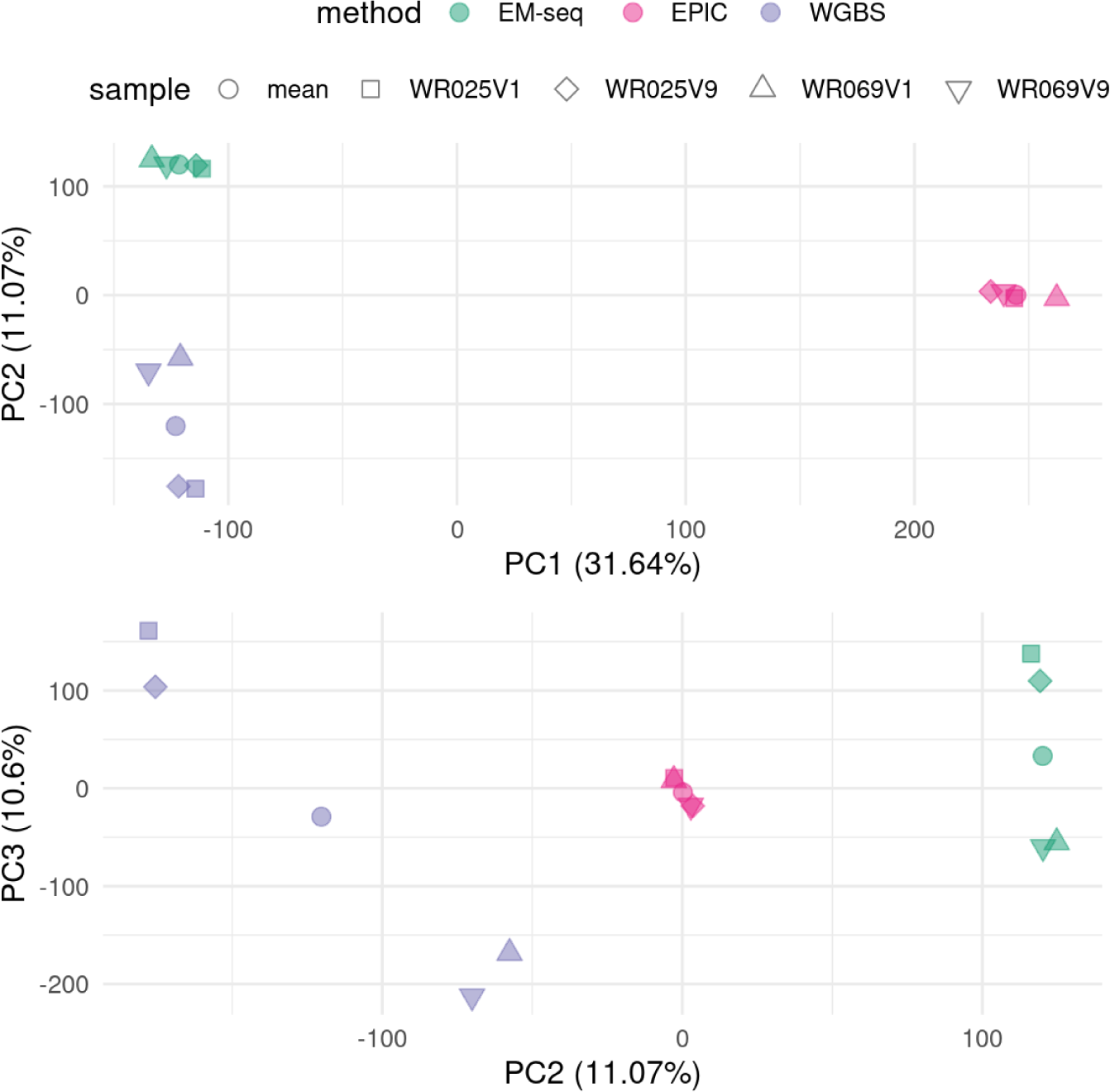
Principal components analysis of per-base methylation levels of four samples across EM-seq, WGBS and EPIC. Variation in the first two principal axes were mainly driven by methodological differences: EPIC and short-read methods occupying separate ends of the first principal component, while the short-read methods separate well on the second principal component. The third principal component (with explained variance value just under the second component) separated by biological sample origin: the four-sided shapes (WR025) and the three-sided ones (WR069) tend to occupy opposite ends. This separation was obvious for the short-read methods but not for EPIC, in line with the lowest variance amongst the EPIC datasets. WGBS had the highest variance. To facilitate downstream comparisons, per-position, per-method mean methylation levels were calculated (filled round points) and included in the plot to confirm they are located amongst the four constituent replicates.

**Supplementary Figure S4.**
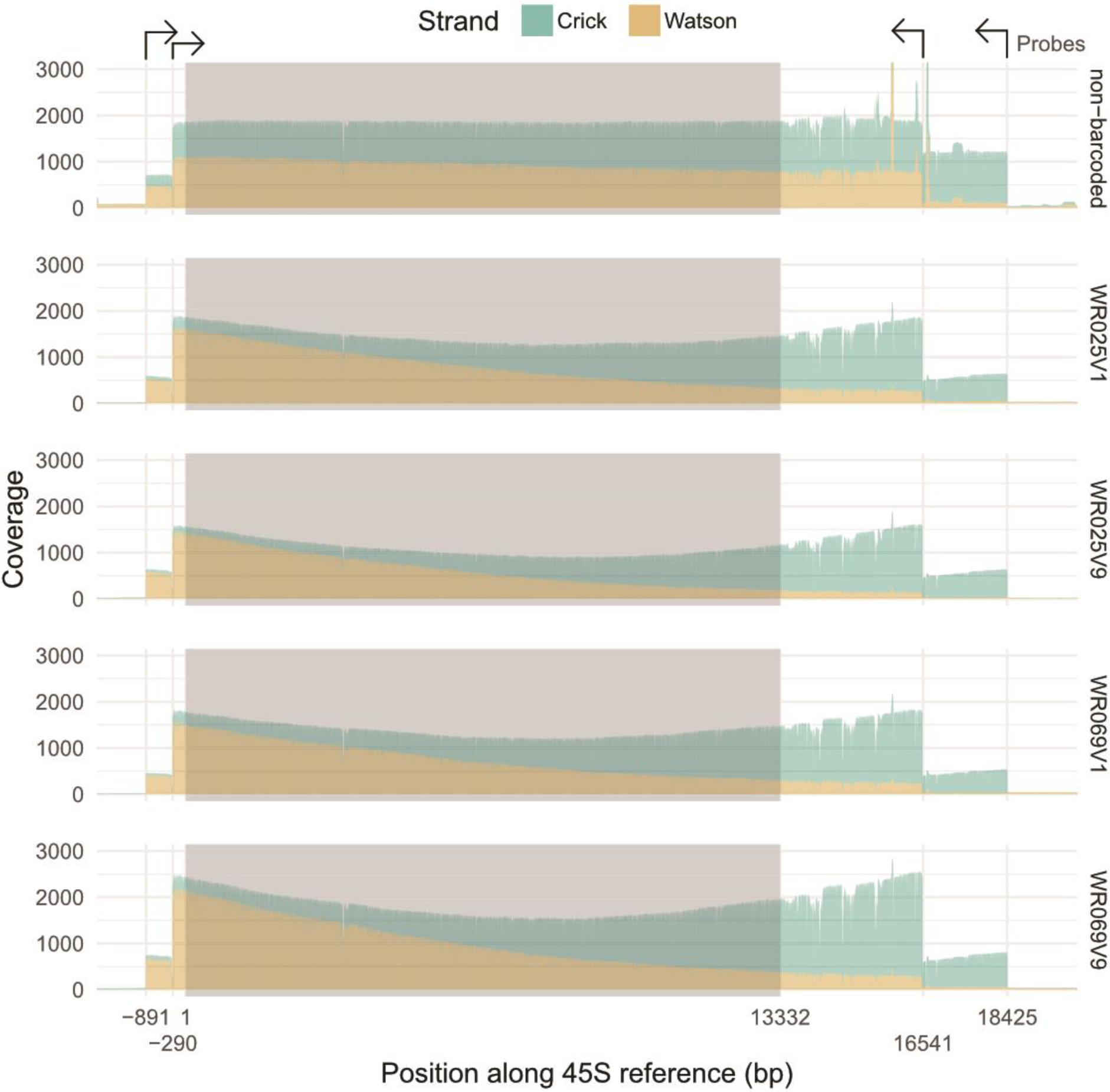
Coverage of mapped reads along the 45S reference used in this experiment. Coverages were plotted separately for each sample (and for reads that were not barcoded). Shaded region represents the transcribed 45S regions (1–13,332 bp). The x-axis positions of -891, -290, 16,451 and 18,425 bp represent the 3’ positions of the probes used in this experiment, and the direction of the probes are indicated at the top of the plot. As Cas9 makes targeted blunt-end cuts in the DNA, the length of the read can be shorter than the gap between cut sites if the other end was sheared/degraded through normal means. This results in a typical U-shaped coverage plot, seen in all barcoded samples. The flatter profile for non-barcoded reads is not well understood (ONT, personal communication).

**Supplementary Figure S5.**
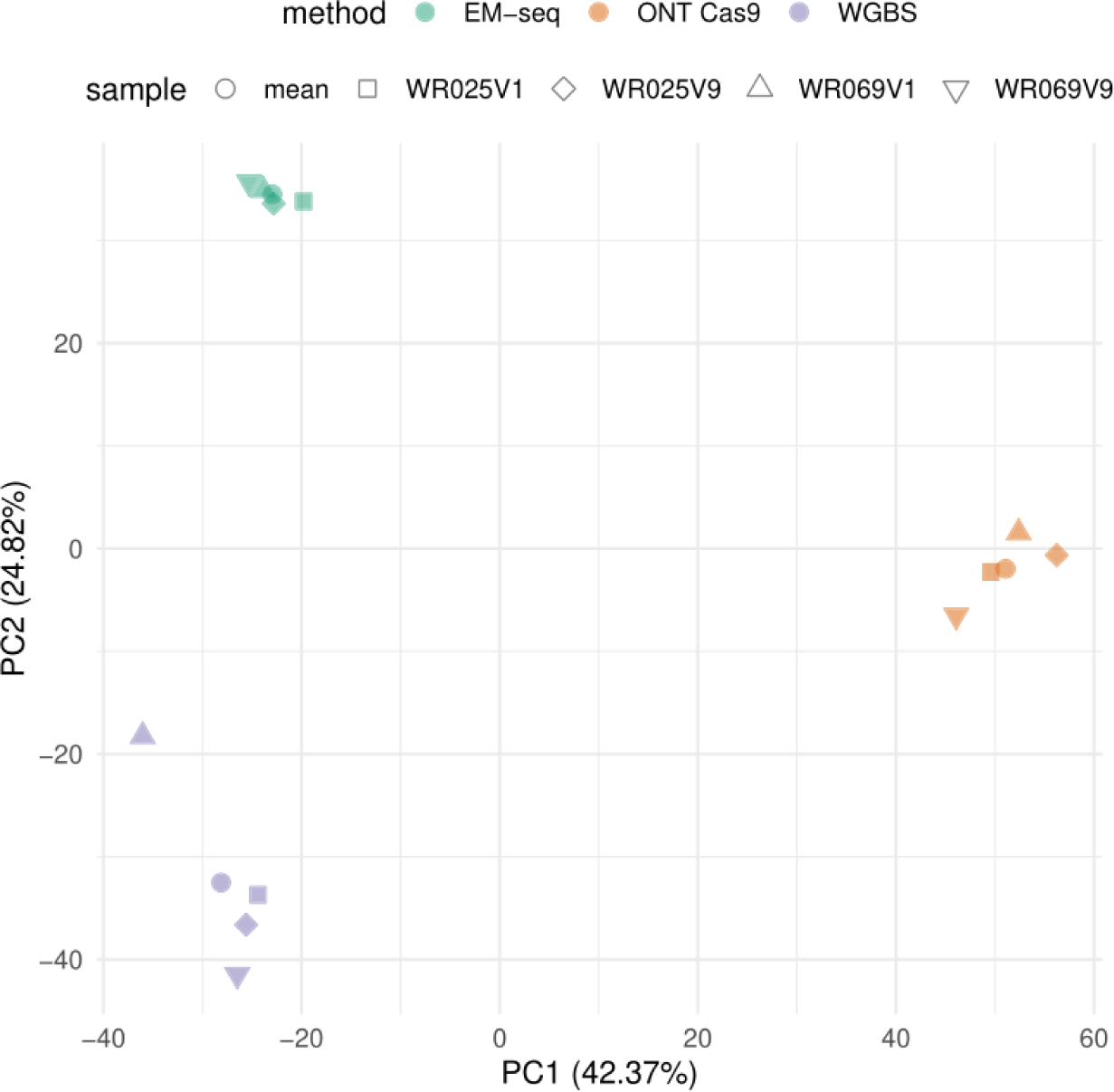
Principal components analysis of per-base methylation levels of four samples across EM-seq, WGBS and ONT Cas9. Variation in the first two principal axes were mainly driven by methodological differences. Replicates within EM-seq datasets had the lowest variation; WGBS highest. To facilitate downstream comparisons, per-position, per-method mean methylation levels were calculated (filled round points) and included in the plot to confirm they are located amongst the four constituent replicates.

**Supplementary Figure S6.**
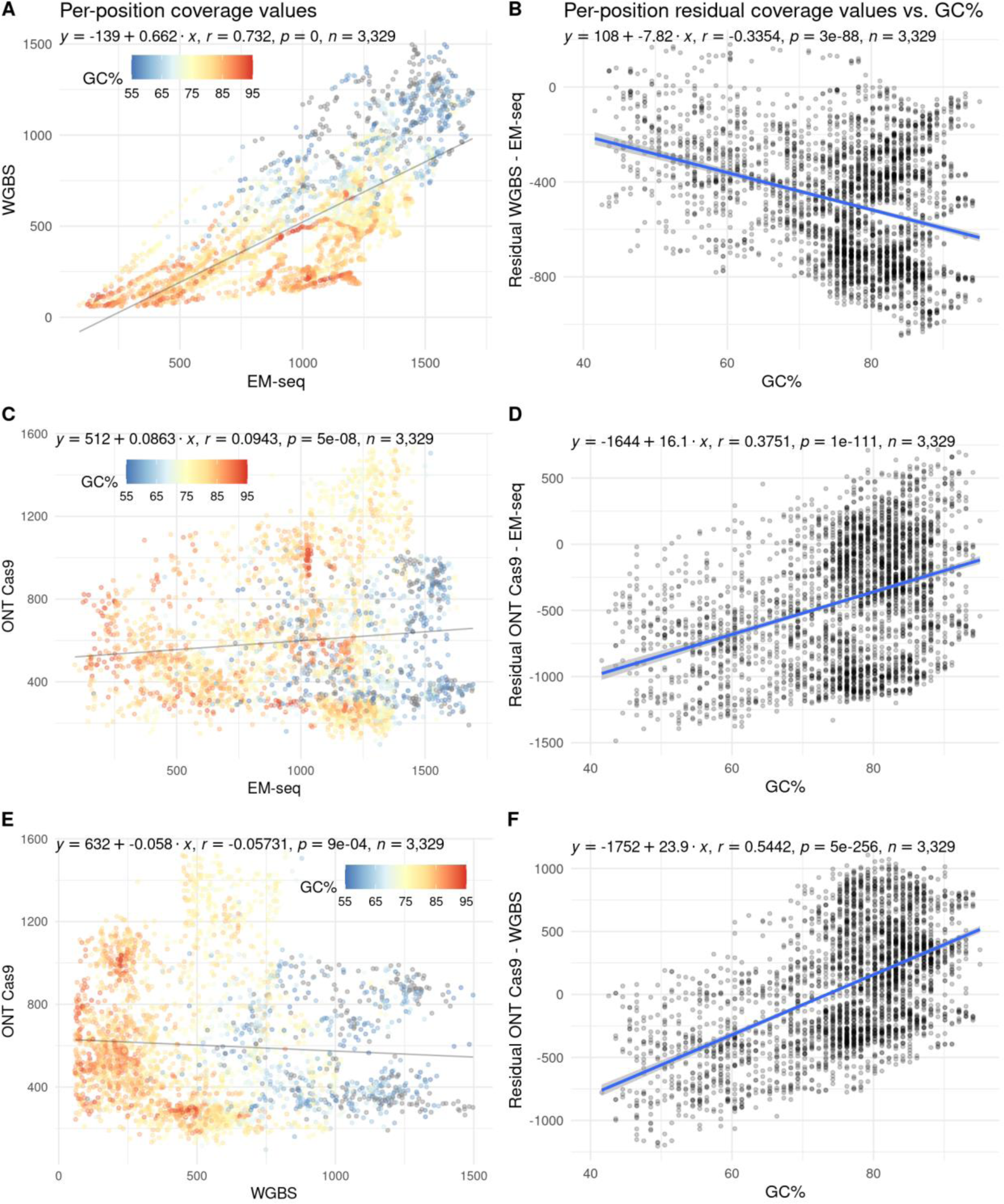
Pairwise comparisons of per-position coverage values from EM-seq, WGBS and ONT Cas9. **A** and **B** are for EM-seq vs. WGBS; **C** and **D** are for ONT Cas9 vs. EM-seq; while **E** and **F** are for ONT Cas9 vs. WGBS. Each point in the plots on the left (**A**, **C** and **E**) represent a single cytosine, and its position on the plot corresponds to the mean coverage value assayed using the method labelled on the axes. It is further coloured by the GC% of its local sequence context (±50 bp) to demonstrate context-dependent biases in coverages. For each point, the residual (differences in coverage values) was computed and plotted against the same GC% value on the right (**B**, **D** and **F**). Coverage values between the short-read methods were better correlated (*r^2^* = 0.54) than either method against ONT Cas9 (*r^2^* < 0.01). EM-seq outperforms WGBS in all available GC% contexts, especially at high GC% contexts (> 75%).

## References

1. Hotchkiss, R.D., The quantitative separation of purines, pyrimidines, and nucleosides by paper chromatography. J Biol Chem, 1948. 175(1): p. 315–32.

2. Doskocil, J. and F. Sorm, Distribution of 5-methylcytosine in pyrimidine sequences of deoxyribonucleic acids. Biochim Biophys Acta, 1962. 55: p. 953–9.

3. Jaenisch, R. and A. Bird, Epigenetic regulation of gene expression: how the genome integrates intrinsic and environmental signals. Nat Genet, 2003. 33 Suppl: p. 245-54.

4. Compere, S.J. and R.D. Palmiter, DNA methylation controls the inducibility of the mouse metallothionein-I gene lymphoid cells. Cell, 1981. 25(1): p. 233–40.

5. Wang, R.Y., C.W. Gehrke, and M. Ehrlich, Comparison of bisulfite modification of 5-methyldeoxycytidine and deoxycytidine residues. Nucleic Acids Res, 1980. 8(20): p. 4777–90.

6. Frommer, M., et al., A genomic sequencing protocol that yields a positive display of 5-methylcytosine residues in individual DNA strands. Proc Natl Acad Sci U S A, 1992. 89(5): p. 1827–31.

7. Harrison, A. and A. Parle-McDermott, DNA methylation: a timeline of methods and applications. Front Genet, 2011. 2: p. 74.

8. Jaksik, R., et al., Microarray experiments and factors which affect their reliability. Biol Direct, 2015. 10: p. 46.

9. Meissner, A., et al., Genome-scale DNA methylation maps of pluripotent and differentiated cells. Nature, 2008. 454(7205): p. 766-70.

10. Lister, R., et al., Highly integrated single-base resolution maps of the epigenome in Arabidopsis. Cell, 2008. 133(3): p. 523–36.

11. Lister, R., et al., Human DNA methylomes at base resolution show widespread epigenomic differences. Nature, 2009. 462(7271): p. 315-22.

12. Pan, H., et al., Measuring the methylome in clinical samples: improved processing of the Infinium Human Methylation450 BeadChip Array. Epigenetics, 2012. 7(10): p. 1173–87.

13. Varinli, H., et al., COBRA-Seq: Sensitive and Quantitative Methylome Profiling. Genes (Basel), 2015. 6(4): p. 1140–63.

14. Pidsley, R., et al., Critical evaluation of the Illumina MethylationEPIC BeadChip microarray for whole-genome DNA methylation profiling. Genome Biol, 2016. 17(1): p. 208.

15. Stirzaker, C., et al., Mining cancer methylomes: prospects and challenges. Trends Genet, 2014. 30(2): p. 75–84.

16. Tanaka, K. and A. Okamoto, Degradation of DNA by bisulfite treatment. Bioorg Med Chem Lett, 2007. 17(7): p. 1912–5.

17. Feng, S., et al., Efficient and accurate determination of genome-wide DNA methylation patterns in Arabidopsis thaliana with enzymatic methyl sequencing. Epigenetics Chromatin, 2020. 13(1): p. 42.

18. Vaisvila, R., et al., Enzymatic methyl sequencing detects DNA methylation at single-base resolution from picograms of DNA. Genome Res, 2021. 31(7): p. 1280–9.

19. Olova, N., et al., Comparison of whole-genome bisulfite sequencing library preparation strategies identifies sources of biases affecting DNA methylation data. Genome Biol, 2018. 19(1): p. 33.

20. Long, M.D., D.J. Smiraglia, and M.J. Campbell, The Genomic Impact of DNA CpG Methylation on Gene Expression; Relationships in Prostate Cancer. Biomolecules, 2017. 7(1).

21. Shi, H., M.X. Wang, and C.W. Caldwell, CpG islands: their potential as biomarkers for cancer. Expert Rev Mol Diagn, 2007. 7(5): p. 519–31.

22. Locke, W.J., et al., DNA Methylation Cancer Biomarkers: Translation to the Clinic. Front Genet, 2019. 10: p. 1150.

23. Laszlo, A.H., et al., Detection and mapping of 5-methylcytosine and 5-hydroxymethylcytosine with nanopore MspA. Proc Natl Acad Sci U S A, 2013. 110(47): p. 18904–9.

24. Schreiber, J., et al., Error rates for nanopore discrimination among cytosine, methylcytosine, and hydroxymethylcytosine along individual DNA strands. Proc Natl Acad Sci U S A, 2013. 110(47): p. 18910–5.

25. Sakamoto, Y., et al., Long-read whole-genome methylation patterning using enzymatic base conversion and nanopore sequencing. Nucleic Acids Res, 2021.

26. Liu, Y., et al., DNA methylation-calling tools for Oxford Nanopore sequencing: a survey and human epigenome-wide evaluation. Genome Biol, 2021. 22(1): p. 295.

27. Yuen, Z.W., et al., Systematic benchmarking of tools for CpG methylation detection from nanopore sequencing. Nat Commun, 2021. 12(1): p. 3438.

28. Delahaye, C. and J. Nicolas, Sequencing DNA with nanopores: Troubles and biases. PLoS One, 2021. 16(10): p. e0257521.

29. Spealman,, P., J. Burrell, and D. Gresham, Inverted duplicate DNA sequences increase translocation rates through sequencing nanopores resulting in reduced base calling accuracy. Nucleic Acids Res, 2020. 48(9): p. 4940–4945.

30. Browne, P.D., et al., GC bias affects genomic and metagenomic reconstructions, underrepresenting GC-poor organisms. Gigascience, 2020. 9(2).

31. Bowen, J., et al., Randomized Trial of a High Protein, Partial Meal Replacement Program with or without Alternate Day Fasting: Similar Effects on Weight Loss, Retention Status, Nutritional, Metabolic, and Behavioral Outcomes. Nutrients, 2018. 10(9).

32. Kulis, M., et al., Whole-genome fingerprint of the DNA methylome during human B cell differentiation. Nat Genet, 2015. 47(7): p. 746–56.

33. Adam, S., et al., Flanking sequences influence the activity of TET1 and TET2 methylcytosine dioxygenases and affect genomic 5hmC patterns. Commun Biol, 2022. 5(1): p. 92.

34. Ravichandran, M., et al., Pronounced sequence specificity of the TET enzyme catalytic domain guides its cellular function. Sci Adv, 2022. 8(36): p. eabm2427.

35. Foox, J., et al., The SEQC2 epigenomics quality control (EpiQC) study. Genome Biol, 2021. 22(1): p. 332.

36. Schröder, C., B. Horsthemke, and C. Depienne, GC-rich repeat expansions: associated disorders and mechanisms. Medizinische Genetik, 2021. 33(4): p. 325–335.

37. Peters, T.J., et al., Evaluation of cross-platform and interlaboratory concordance via consensus modelling of genomic measurements. Bioinformatics, 2019. 35(4): p. 560–570.

38. Jain, S., et al., Methylation of the CpG sites only on the sense strand of the APC gene is specific for hepatocellular carcinoma. PLoS One, 2011. 6(11): p. e26799.

39. Chen, Y.A., et al., Discovery of cross-reactive probes and polymorphic CpGs in the Illumina Infinium HumanMethylation450 microarray. Epigenetics, 2013. 8(2): p. 203–9.

40. Gilpatrick, T., et al., Targeted nanopore sequencing with Cas9-guided adapter ligation. Nat Biotechnol, 2020. 38(4): p. 433–438.

41. Payne, A., et al., Readfish enables targeted nanopore sequencing of gigabase-sized genomes. Nat Biotechnol, 2021. 39(4): p. 442–450.

42. Krueger, F. and S.R. Andrews, Bismark: a flexible aligner and methylation caller for Bisulfite-Seq applications. Bioinformatics, 2011. 27(11): p. 1571–2.

43. Kim, J.H., et al., Variation in human chromosome 21 ribosomal RNA genes characterized by TAR cloning and long-read sequencing. Nucleic Acids Res, 2018. 46(13): p. 6712–6725.

44. Gonzalez, I.L. and J.E. Sylvester, Complete sequence of the 43-kb human ribosomal DNA repeat: analysis of the intergenic spacer. Genomics, 1995. 27(2): p. 320–8.

45. Parks, M.M., et al., Variant ribosomal RNA alleles are conserved and exhibit tissue-specific expression. Sci Adv, 2018. 4(2): p. eaao0665.

46. Hansen, K.D., B. Langmead, and R.A. Irizarry, BSmooth: from whole genome bisulfite sequencing reads to differentially methylated regions. Genome Biol, 2012. 13(10): p. R83.

47. Wu, H., C. Wang, and Z. Wu, A new shrinkage estimator for dispersion improves differential expression detection in RNA-seq data. Biostatistics, 2013. 14(2): p. 232–43.

48. Wickham, H., ggplot2: Elegant Graphics for Data Analysis. 2016: Springer-Verlag New York.

49. Gu, Z., R. Eils, and M. Schlesner, Complex heatmaps reveal patterns and correlations in multidimensional genomic data. Bioinformatics, 2016. 32(18): p. 2847–9.

50. Ou, J., et al., motifStack for the analysis of transcription factor binding site evolution. Nat Methods, 2018. 15(1): p. 8–9.

51. Triche T.J.,, Jr., et al., Low-level processing of Illumina Infinium DNA Methylation BeadArrays. Nucleic Acids Res, 2013. 41(7): p. e90.

52. Ziller, M.J., et al., Coverage recommendations for methylation analysis by whole-genome bisulfite sequencing. Nat Methods, 2015. 12(3): p. 230–2, 1 p following 232.

53. Labun, K., et al., CHOPCHOP v3: expanding the CRISPR web toolbox beyond genome editing. Nucleic Acids Res, 2019. 47(W1): p. W171–w174.

